# The role of antigen recognition in the γδ T cell response at the controlled stage of *M. tuberculosis* infection

**DOI:** 10.1101/2021.09.14.460324

**Authors:** Roshni Roy Chowdhury, John R. Valainis, Oliver Kask, Mane Ohanyan, Meng Sun, Huang Huang, Megha Dubey, Lotta von Boehmer, Elsa Sola, Xianxi Huang, Patricia K. Nguyen, Thomas J. Scriba, Mark M. Davis, Sean C. Bendall, Yueh-hsiu Chien

**Affiliations:** Department of Microbiology and Immunology, Stanford University, Stanford, CA, USA; Program in Immunology, Stanford University, Stanford, CA, USA; Department of Pathology, Stanford University, Stanford, CA, USA; Institute for Immunity, Transplantation, and Infection, Stanford University, Stanford, CA, USA; Division of Cardiovascular Medicine, Stanford University, Stanford, CA, USA; The First Affiliated Hospital of Shantou University Medical College, Shantou, China; South African Tuberculosis Vaccine Initiative, Institute of Infectious Disease and Molecular Medicine and Division of Immunology, Department of Pathology, University of Cape Town, Cape Town, South Africa; The Howard Hughes Medical Institute, Stanford University School of Medicine, Stanford, CA, USA

## Abstract

*γδ* T cells contribute to host immune defense uniquely; but how they function in different stages (e.g., acute versus chronic) of a specific infection remains unclear. As the role of *γδ* T cells in early, active *Mycobacterium tuberculosis* (Mtb) infection is well documented, we focused on elucidating the *γδ* T cell response in persistent or controlled Mtb infection. Systems analysis of circulating *γδ* T cells from a South African adolescent cohort identified a distinct population of CD8+ *γδ* T cells that expanded in this state. These cells had features indicative of persistent antigenic exposure but were robust cytolytic effectors and cytokine/chemokine producers. While these *γδ* T cells displayed an attenuated response to TCR-mediated stimulation, they expressed Natural Killer (NK) cell receptors and had robust CD16 (Fc*γ*RIIIA)-mediated cytotoxic response, suggesting alternative ways for *γδ* T cells to control this stage of the infection. Despite this NK-like functionality, the CD8+ *γδ* T cells consisted of highly expanded clones, which utilized TCRs with different V*γ*/*δ* pairs. Theses TCRs could respond to an Mtb-lysate, but not to phosphoantigens, which are components of Mtb-lysate that activate *γδ* T cells in acute Mtb infection, indicating that the CD8+ *γδ* T cells were induced in a stage-specific, antigen-driven manner. Indeed, trajectory analysis showed that these *γδ* T cells arose from naive cells that had traversed distinct differentiation paths in this infection stage. Importantly, increased levels of CD8+ *γδ* T cells were also found in other chronic inflammatory conditions, including cardiovascular disease and cancer, suggesting that persistent antigenic exposure may lead to similar *γδ* T cell responses.

## Introduction

A better understanding of how lymphocytes control both the acute and chronic phase of the same infection is key to the development of new intervention strategies and improved vaccines. The involvement of *γδ* T cells in several acute and chronic infections has been noted for some time. In particular, in individuals with active tuberculosis (TB) disease, and other acute bacterial and parasitic infections, *γδ* T cell frequencies can increase in the peripheral blood from <5% in healthy subjects to >45% in certain patients^1^. These *γδ* T cells primarily express V*γ*9V*δ*2 TCRs, which respond to phosphoantigens – metabolites generated in the isoprenoid pathway in bacteria and parasites. Activated *γδ* T cells can eliminate infected cells or pathogens *in vitro*^2–5^, suggesting that direct killing by these cells is part of how the infection is controlled. More recently, increases in the frequency of circulating IL-17+ V*γ*9V*δ*2 T cells were observed in children with acute bacterial meningitis^6^ and in patients with active pulmonary tuberculosis^7^, consistent with the findings in mouse models of infection, where *γδ* T cells are the major initial IL-17 producers in acute infection that initiate the inflammatory response^8^.

*γδ* T cells have also been implicated in controlling chronic viral infections^9–11^. Selective *γδ* T cell populations expand in the peripheral blood of patients infected with HIV, CMV, EBV, or HSV^12–14^. In immunocompromised transplant patients with CMV reactivation, the rise of circulating *γδ* T cells correlates with the resolution of infection, and increased numbers of *γδ* T cells are also associated with improved long-term survival in these patients with high risk of CMV reactivation and tumor development^15^. In these cases, most of the activated *γδ* T cells express TCRs encoded by V*δ*1 and V*δ*3, paired with different V*γ* chains, some of which show reactivity to virus-infected cells and tumor cell lines^15^.

While these studies highlight the ability of *γδ* T cells to respond to pathogenic challenges in a context-dependent manner, it is unclear whether the difference in these responses reflect different pathogenic challenges (bacteria or parasites vs. viruses) and/or the chronicity of the infection. This question is particularly relevant to Mtb infection, where most infections manifest as a clinically asymptomatic state, presumably held in check, or cleared by the host immune system. This state, previously known as latent TB, and referred to as controlled Mtb infection here, may be best defined as a state of persistent immune response to Mtb antigens detected by an interferon *γ* (IFN*γ*)-release assay, but without signs or symptoms of active disease. Previously, we have characterized the immune state associated with this controlled infection stage in a cohort of South African adolescents aged 13–18 years^16^. This cohort is from a highly endemic area but has a lower rate of active TB disease than is seen in young children and adults^17^, indicating a well-controlled Mtb infection. Our results indicated that *γδ* T cells contribute to the concerted effort toward Mtb infection control. However, the *γδ* T cell response in these subjects remains largely uncharacterized.

Here, we employed mass cytometric analysis, transcriptomic profiling, and functional assays, together with a newly developed Cytoskel algorithm to construct pseudotime trajectories, which reflect cellular progression through differentiation stages to study *γδ* T cells from peripheral blood mononuclear cells (PBMCs) from the same South African adolescent cohort. Our results identify unique features of *γδ* T cells in this stage of the infection and underscore the importance of antigen recognition in this phase of the *γδ* T cell response.

## Results

### An increased frequency of peripheral γδ T cells with robust effector functions and features of persistent antigenic exposure is associated with controlled Mtb infection

To identify the *γδ* T cell subpopulation(s) that associate with controlled Mtb infection, we analyzed our previously generated high-dimensional cytometry by time-of-flight (CyTOF) datasets, which examined cell surface markers, intracellular cytokines/cytolytic effectors, and signaling capacity (the ability to transiently phosphorylate signaling effectors in response to stimulation) from PBMCs of uninfected donors and individuals with controlled Mtb infection from the South African adolescent cohort^16^. First, we used the Citrus (cluster identification, characterization, and regression) algorithm^18^ to identify *γδ* T cell populations that stratify individuals with controlled Mtb infection versus uninfected donors based on cell surface marker expression and intracellular cytokines/cytolytic staining. This unbiased and unsupervised analysis revealed that several *γδ* T cell clusters (Clusters 1-5), which shared common features, including the expression of CD8, were present at significantly higher frequencies (False Discovery Rate (FDR) < 0.01) in individuals with controlled infection compared to uninfected donors (N=14/group). In addition, there was a significant decrease in the frequency of the *γδ* T cell subpopulation which did not express CD8 (Cluster 6, comprised of the majority of the CD8-*γδ* T cells) (**Extended Data Figs. 1a-e**), reflecting the observation that the total *γδ* T cell frequencies in circulation remain similar between the two groups^16^. We confirmed this observation by FACS analysis of an additional 17 uninfected donors and 19 individuals with controlled Mtb infection (**Fig. 1a**).

**Figure 1.**
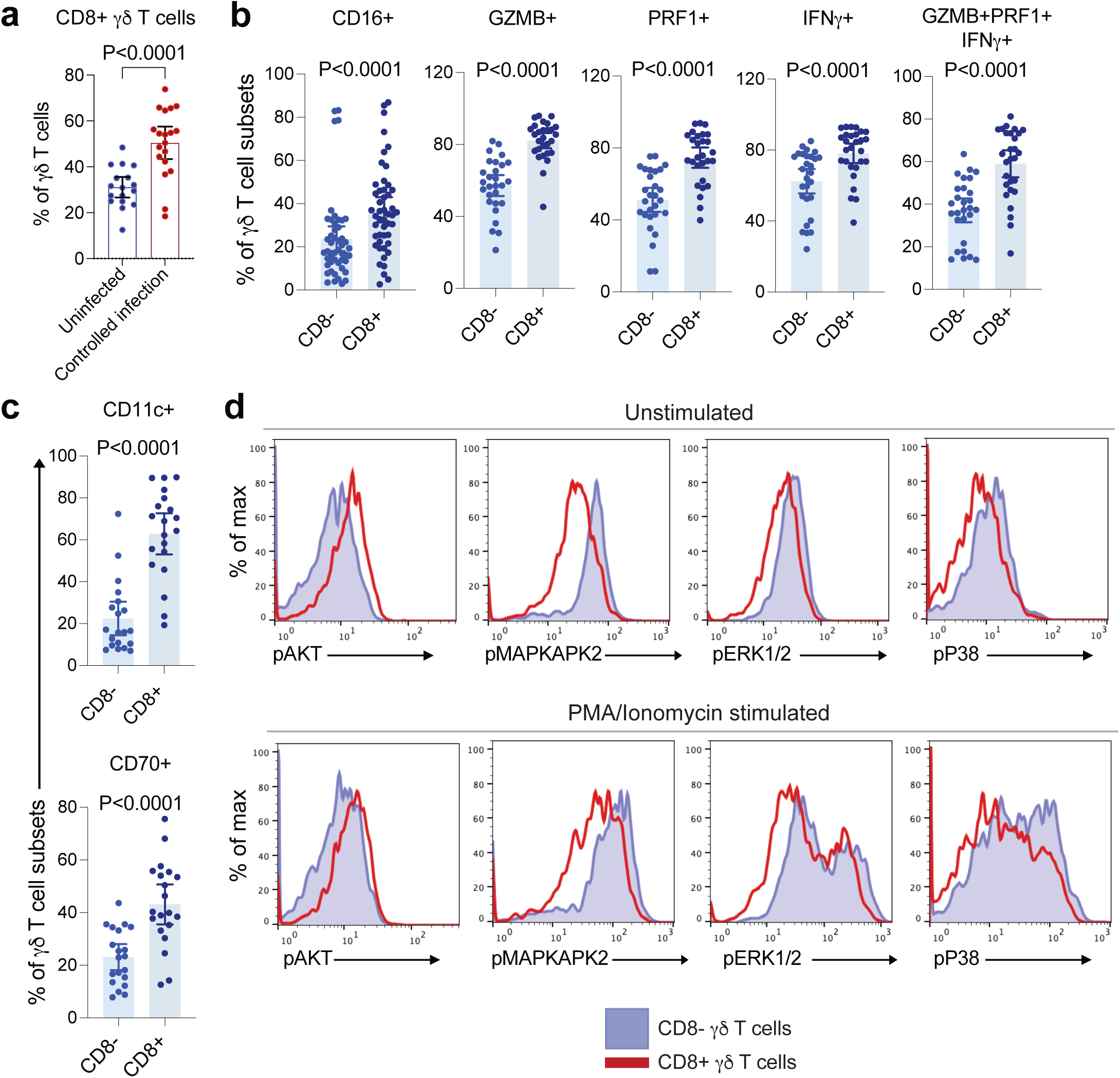
An Increased frequency of periphery CD8+ γδ T cells with effector phenotype associates with controlled Mtb infection. **(a)** Percentage of circulating CD8+ *γδ* T cells in 17 uninfected donors and 19 donors with controlled Mtb infection from a South African adolescent cohort. *P*-value was derived using the Mann-Whitney test. Error bars represent mean and 95% confidence intervals. **(b, c)** Percentages of CD16+, GZMB+, PRF1+, IFN*γ*+, and polyfunctional (GZMB+PRF1+IFN*γ*+) cells (b), and CD11c+ and CD70+ cells (c), within CD8- and CD8+ *γδ* T cell subsets. *P*-values were derived using Wilcoxon matched-pairs signed rank test. Error bars represent mean and 95% confidence intervals. **(d)** Representative histograms showing the phosphorylation levels of signaling effectors in CD8- and CD8+ *γδ* T cell subsets at baseline (top panel) and post PMA-Ionomycin stimulation (bottom panel).

viSNE dimensionality reduction of the CyTOF analysis showed that the CD8+ *γδ* T cells clustered together and displayed features of terminally differentiated effectors, which regain CD45RA expression (CD45RA+CD45RO-CCR7-CD27-) (**Extended Data Figs. 2a, b**). These cells had a high expression of cytotoxic molecules (GZMB+PRF1+) and significantly higher proportions of CD16+ cells relative to the CD8- *γδ* T cell subpopulation (**Fig. 1b, Extended Data Figs. 2a, b**), suggesting enhanced cytolytic ability, which could be mediated through CD16. The CD8+ subset also comprised of significantly higher frequencies of GZMB+, PRF1+, and IFN*γ*+ cells, as well as polyfunctional cells (**Fig. 1b**). Interestingly, these CD8+ *γδ* T cells were distinguished by their significantly increased expression of two classical myeloid cell markers, CD11c and CD70 (**Fig. 1c, Extended Data Fig. 2a**), whose expression on *αβ* T cells has been linked to antigen-driven responses and chronic immune activation^19, 20^. Consistently, CD8+ *γδ* T cells showed small but significantly lower cell surface CD3 expression as compared to CD8- *γδ* T cells (**Extended Data Fig. 1e).** This observation was also confirmed by FACS analysis of the 17 uninfected donors and 19 individuals with controlled Mtb infection (**Extended Data Fig. 2c**).

The CD8+ *γδ* T cells also differed in terms of their activation state and signaling capacity, displaying enhanced AKT phosphorylation at baseline (unstimulated condition) and post stimulation with phorbol myristate acetate (PMA) and ionomycin in comparison to CD8- *γδ* T cells (**Fig. 1d, Extended Data Fig. 2d).** In cytotoxic CD8+ *αβ* T cells, AKT activation has been shown to play an important role in driving TCR- and IL-2-induced transcriptional programs that control the expression of cytokines/chemokines and cytolytic effectors, regulate trafficking responses, and cell fate determination^21^. Relative to CD8- *γδ* T cells, the CD8+ *γδ* T cell subset displayed significantly lower phosphorylation levels of the mitogen-activated protein kinases (MAPKs), including ERK-1/2, p38, and MAPKAPK2, a substrate for p38, at baseline and upon stimulation with PMA and ionomycin (**Fig. 1d, Extended Data Fig. 2d**). Defects in the MAPK pathway and induction of ERK phosphorylation have been associated with reduced proliferation^22^. These observations are consistent with these cells being terminally differentiated effectors.

To broaden our analysis, we performed bulk RNA sequencing (bulk-RNA-seq) on FACS sorted CD8+ and CD8- *γδ* T cells from peripheral blood samples of five donors with controlled Mtb infection and identified 1,273 significantly (Adjusted *P*<0.05) differentially expressed genes (**Fig. 2a, Supplementary Table 1**). Compared to CD8- *γδ* T cells, the CD8+ *γδ* T cells showed significantly higher levels of some genes associated with prior antigenic exposure and chronic TCR activation in *αβ* T cells, such as TIGIT, LAG3, CD244 [2B4], B3GAT1 [CD57], PRDM1 [Blimp-1], NR4A2, and TOX (**Fig. 2b**), and displayed cytolytic features characterized by high expression of GNLY (Granulysin), PRF-1 (Perforin-1), granzymes (GZMA, GZMH), and NKG7 (a regulator of exocytosis of cytotoxic granules) (**Fig. 2b, Supplementary Table 1**). The CD8+ *γδ* T cells also expressed higher levels of both FCGR3A (CD16a) (did not reach statistical significance) and FCGR3B (CD16b) (**Fig. 2b, Supplementary Table 2**). Furthermore, they showed a preferential upregulation of NK-cell-associated activating and inhibitory receptor genes, including KLRD1(CD94), KLRC3 (NKG2E), KLRC2 (NKG2C), KLRG1, KLRC4 (NKG2F), NCR1 (NKp46), CD244 (2B4), and the CD158 family of KIR receptors: KIR2DS4, KIR2DL1, KIR2DL3, KIR3DL1, KIR3DL2, and KIR3DL3 (**Fig. 2b, Supplementary Table 2**). Although not statistically significant, other NK-related genes like KLRF1 (NKp80), and KLRK1 (NKG2D) also showed higher expression levels in CD8+ compared to CD8- *γδ* T cells (**Supplementary Table 2**). Taken together, these findings were consistent with the CyTOF analysis, which indicated that CD8+ *γδ* T cells are highly cytolytic effectors and that their response may be modulated by NK cell receptors. These cells may also deploy NK cell-like functions through CD16.

**Figure 2.**
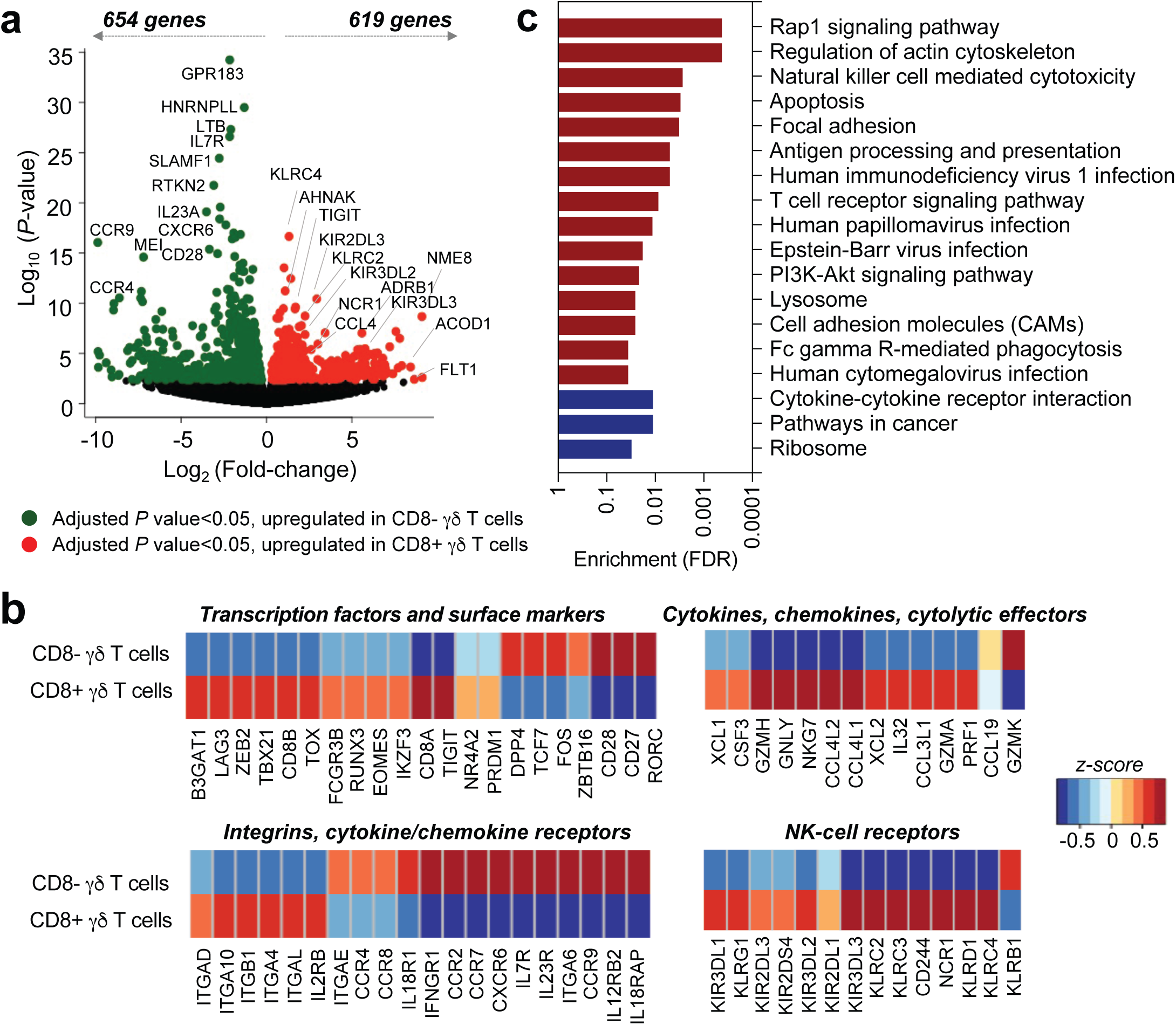
Peripheral CD8+ γδ T cells exhibit NK cell like features and characteristics indicative of prior antigenic exposure. **(a)** Volcano plot showing differential gene expression between bulk sorted CD8- and CD8+ *γδ* T cells derived from peripheral blood samples of 5 donors with controlled Mtb infection. **(b)** Heatmaps showing the expression patterns of selected genes significantly differentially expressed (adjusted *P* <0.05) between CD8- and CD8+ *γδ* T cells. **(c)** KEGG Pathway analysis of transcripts differentially upregulated in CD8+ *γδ* T cells (red bars) and CD8- *γδ* T cells (blue bars) analyzed by the ShinyGO gene-set enrichment tool, which calculates enrichment based on hypergeometric distribution followed by FDR correction. The pathways listed have enrichment FDR<0.05.

Although the abundance of cytokine mRNAs with a short half-life cannot be detected accurately in gene expression assays without re-stimulation (to re-induce their expressions after cell isolation) as we did here, we found very high levels of IL-32 expression, which has been implicated in various chronic inflammatory diseases^23^. Furthermore, both the CD8+ and the CD8- *γδ* T cell subsets expressed high levels of the chemokine CCL5 (RANTES) (**Supplementary Table 2)**. Nonetheless, the CD8+ *γδ* T cells expressed significantly higher levels of XCL1, XCL2, CSF3, CCL4L1(MIP1b), CCL4L2, CCL3L1 (MIP1AP), and CCL19 (**Fig. 2b**), which target a wide range of immune cells, including lymphocytes, dendritic cells, and other myeloid cells, suggesting that mobilizing leukocytes to the site of the infection may be one of the ways that CD8+ *γδ* T cells regulate the inflammatory response. In fact, XCL1-producing activated CD8+ *αβ* T cells were reported to orchestrate the trafficking of IFN*γ*+ CD4 T cells in a mouse model of chronic tuberculosis infection^24^.

Compared to the CD8+ *γδ* T cells, the CD8- subset showed higher expression of RORC, SOX4, ID3, and LEF1 – transcription factors associated with IL-17-committed *γδ* T cells, but relatively lower production of lytic effector molecules (**Fig. 2b**). While all *γδ* T cells expressed high levels of ITGB2 (LFA-1), a suite of integrin genes – ITGAD (CD11D), ITGA10, ITGAL (CD11A), ITGA4 (CD49D), ITGB1 (VLAB) and the chemokine receptor CX_3_CR1 were preferentially upregulated in CD8+ *γδ* T cells (**Fig. 2b, Supplementary Table 2)**, suggesting specific tissue homing properties to extra-intestinal sites, including the central nervous system, lungs and salivary glands^25^. The CD8- *γδ* T cells, in contrast, expressed higher levels of the gut-homing receptor ITGAE (*α*E).

KEGG (Kyoto Encyclopedia of Genes and Genomes) enrichment pathway analysis showed that relative to CD8- *γδ* T cells, CD8+ *γδ* T cells had a significant enrichment for genes associated with Fc*γ*R-mediated functions, and those encoding components of the TCR, PI3K-AKT, and Rap1 signaling machinery (**Fig. 2c**). Signaling via Rap1, whose activation by TCR ligation and chemokines is known to regulate integrin- (specifically, ITGAL and ITGA4)-mediated cell adhesion^26^, may reflect the mechanistic underpinnings of the migratory behavior of peripheral CD8+ *γδ* T cells. In addition, CD8+ *γδ* T cells had a significant enrichment for genes involved in immune response elements associated with chronic viral infections, such as CMV, EBV, and HIV (**Fig. 2c**), highlighting the similarity between *γδ* T cell responses in controlled Mtb and chronic viral infections.

Taken together, these analyses indicated that CD8+ *γδ* T cells in individuals with controlled Mtb infection were highly cytolytic, multi-functional effectors with altered response to TCR-mediated activation, but with the ability to mount CD16-mediated responses. To test these suppositions, we first assayed the intake of MitoTracker-Green by CD8- and CD8+ *γδ* T cells to compare their mitochondrial mass. Different states of activated *αβ* T cells are known to have different patterns of metabolism, but all effector *αβ* T cells have higher mitochondrial mass when compared with naïve or exhausted T cells^27^. We found that compared to CD8- *γδ* T cells, the CD8+ *γδ* T cells showed significantly higher mitochondrial mass (**Fig. 3a**). To evaluate these cells’ response to TCR-mediated activation, we stimulated PBMCs *in vitro* with either anti-CD3 antibody or an Mtb- lysate and used the upregulation of CD69 as a read-out for T cell activation. We also demonstrated that the response to Mtb-lysate was inhibited by Cyclosporine A, indicating that the activation was mediated through the TCR. We found that both the percentage of cells, which upregulated CD69, and the CD69 expression level, were lower on the CD8+ compared to the CD8- *γδ* T cells (**Fig. 3b**). This indicated that these CD8+ *γδ* T cells were hypo-responsive to antigenic challenge. Nonetheless, by measuring antibody-dependent CD107a degranulation from isolated *γδ* T cells and from total PBMCs, we found that the antibody-dependent cellular cytotoxicity (ADCC) potential of the CD8+ *γδ* T cells was significantly higher than the CD8- *γδ* T cells and was comparable to that of NK cells (**Figs. 3c, d, Extended Data Fig. 2e**).

**Figure 3.**
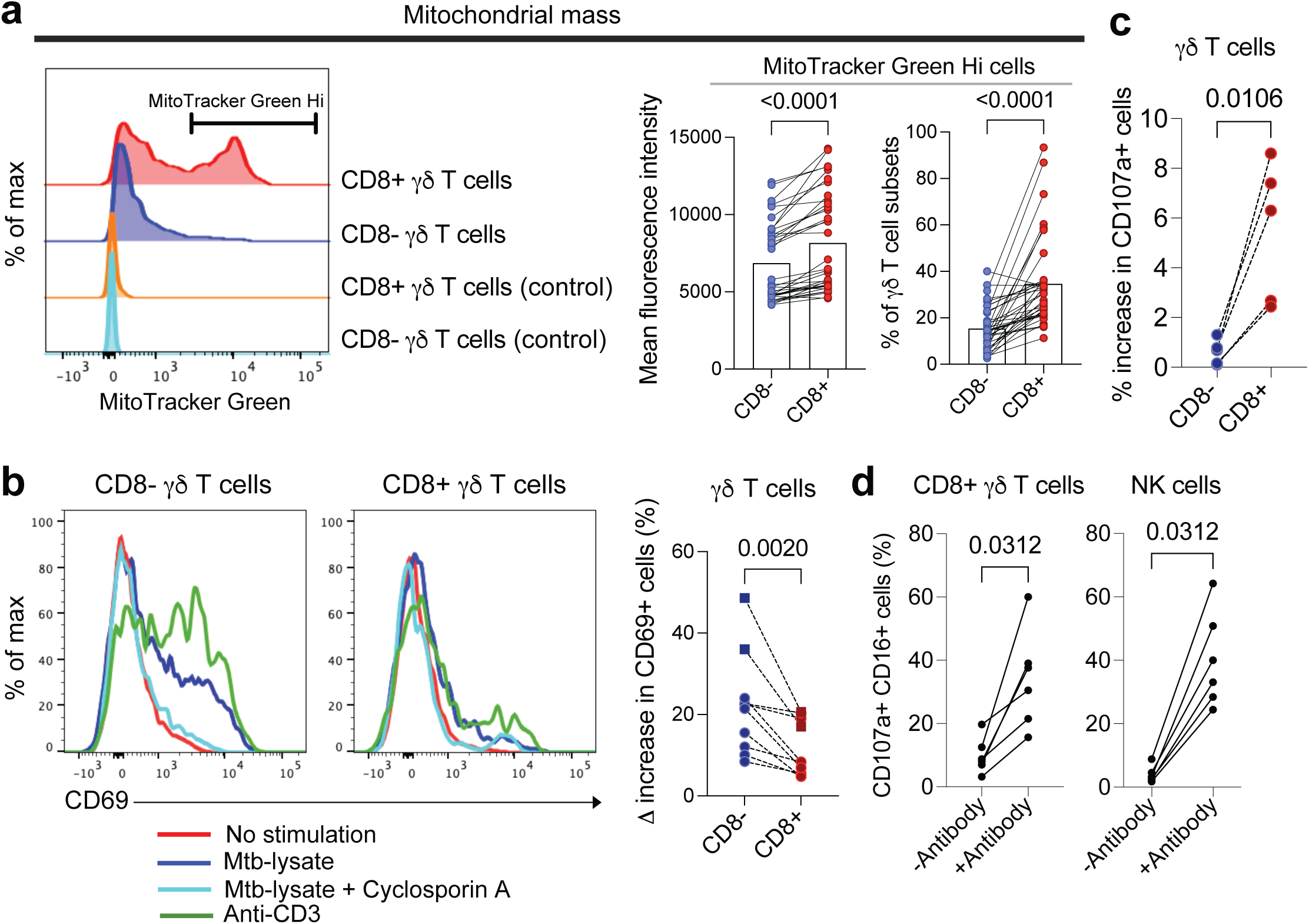
Peripheral CD8+ γδ T cells are effectors which are hyporesponsive to TCR mediated signaling but can mount CD16 mediated ADCC response. **(a)** Histograms showing MitoTracker Green staining of CD8- and CD8+ *γδ* T cells (left). The mean fluorescence intensity (middle) and the percentage of MitoTracker Green Hi high cells (right) within the CD8- and CD8+ *γδ* T cell subsets (right) are shown (N=17 uninfected and 19 controllers). *P*-values were derived using Wilcoxon matched-pairs signed rank test. **(b)** Histograms showing the expression of CD69 on CD8- and CD8+ *γδ* T cells under unstimulated and stimulated conditions (Mtb-lysate, Mtb- lysate with Cyclosporin A, anti-CD3) (left) and the percentage change in CD69+ cells post stimulation in CD8- and CD8+ *γδ* T cells (right). Circles represent Mtb-lysate stimulation, while squares represent anti-CD3 stimulation. *P*-value was derived using Wilcoxon matched-pairs signed rank test. **(c)** ADCC response of CD8- and CD8+ *γδ* T cells, measured by the % increase in CD107a degranulation. Total *γδ* T cells were sorted and incubated in the presence of antibody- coated or uncoated P815 target cells (N=5 donors). *P*-value was derived using Wilcoxon matched-pairs signed rank test. **(d)** ADCC response of CD8+ *γδ* T cells and NK cells measured by incubating total PBMCs in the presence of antibody coated or uncoated P815 target cells (N=6 donors). *P*-values were derived using Wilcoxon matched-pairs signed rank test.

### CD8+ γδ T cells express clonally focused TCRs and respond to Mtb-lysate

Given the observation that the CD8+ *γδ* T cells were hypo-responsive to antigenic challenge, we sought to determine if the increased frequency of peripheral CD8+ *γδ* T cells in controlled Mtb infection reflected antigen-driven responses. To this end, we performed direct *ex vivo* single-cell TCR sequencing (scTCR-seq)^28^ on CD8+ and CD8- *γδ* T cells isolated from peripheral blood samples of four donors with controlled Mtb infection. We found that in all donors, the CD8+ *γδ* TCRs were composed of a few dominant expanded clonotypes (multiplets). In contrast, most of the CD8- *γδ* TCR clonotypes appeared only once (singletons) (**Fig. 4a**). The percent clonality (calculated as the proportion that the multiplets occupy in the total repertoire) across all donors showed a significant difference between the CD8+ and CD8- *γδ* T cells (*P*=0.0045) (**Fig. 4b**). While no identical TCRs in the CD8+ *γδ* TCR repertoire were found between different donors, greater sequencing coverage will be required to determine whether there are shared TCR sequences among individuals. Regardless, these observations suggest that the peripheral CD8+ *γδ* T cells consist of antigen-expanded *γδ* T cell clones. Importantly, while acute Mtb infection induces increased frequencies of circulating *γδ* T cells expressing V*γ*9V*δ*2 TCRs^29^, the repertoire of most circulating CD8+ *γδ* T cells in controlled Mtb infection was skewed towards V*δ*1 and to a lesser extent V*δ*3 gene usage. These TCRs displayed chain-pairing diversity with V*γ*2/3/4 (**Fig. 4c, Supplementary Table 3**). The TCR V gene usage of expanded *γδ* T cells in controlled Mtb infection was similar to those described for *γδ* T cells that expand in chronic CMV infection^30^.

**Figure 4.**
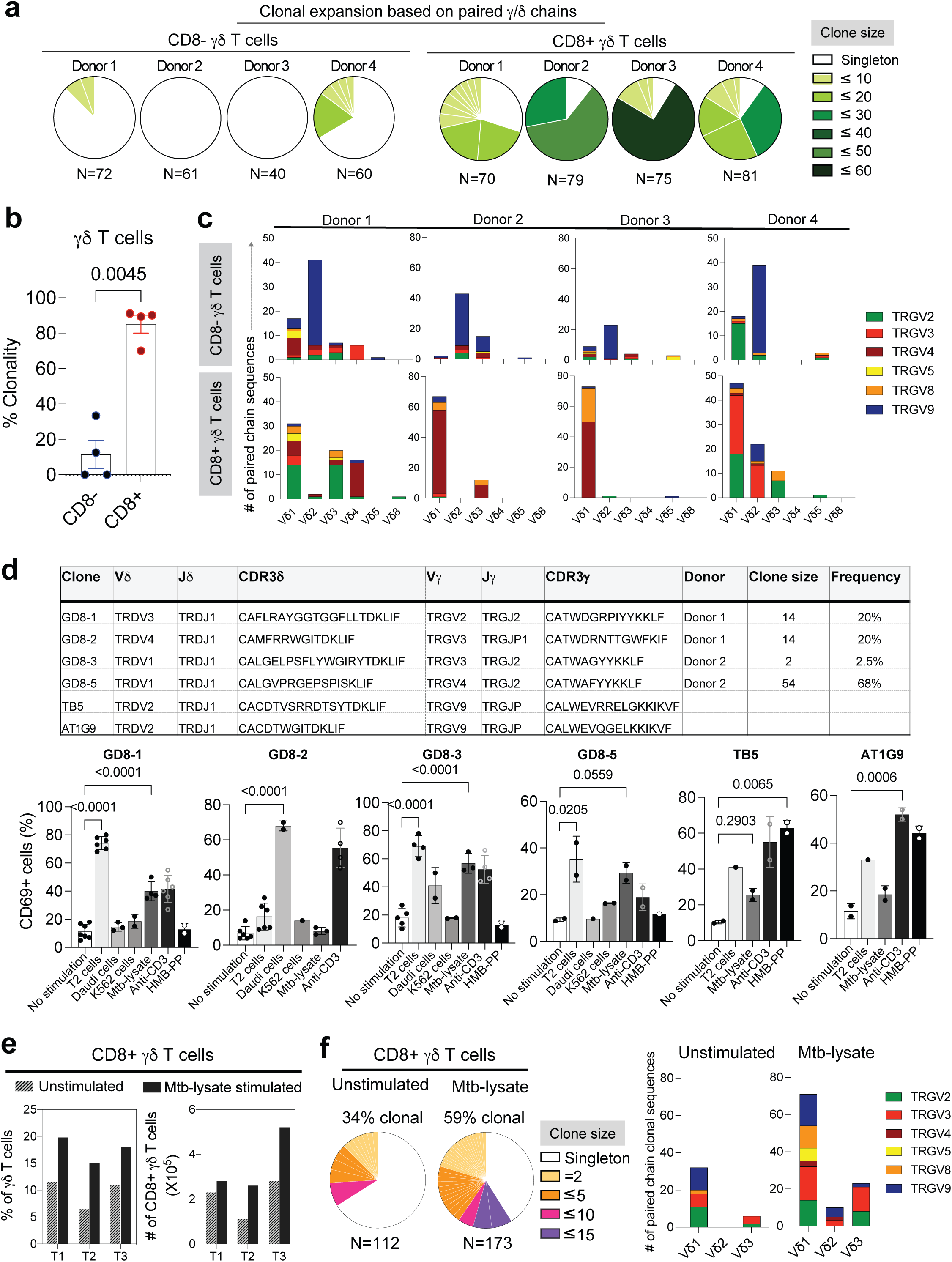
CD8+ γδ T cells express clonally focused TCRs and respond to Mtb-lysate. **(a)** Pie charts depicting the clonal expansion of circulating CD8- and CD8+ *γδ* T cells in 4 donors with controlled Mtb infection. The number of cells with both *γ* and *δ* chains identified is shown below each pie chart. For each TCR sequence expressed by two or more cells (clonally expanded), the absolute number of cells expressing that clone is shown by a distinct colored section. **(b)** Percent clonality of CD8- and CD8+ *γδ* T cells determined from paired single cell *γ* and *δ* chain sequencing. Clonality is defined as the total number of sequences that appear more than once relative to the total number sequenced per sample. *P*-value was determined using the paired *t* test. Error bars represent mean and 95% confidence intervals. **(c)** Stacked bar plots showing the V*γ*/*δ* chain pairings of CD8- (top) and CD8+ (bottom) *γδ* T cells isolated from the 4 donors with controlled Mtb infection. **(d)** CDR3*δ*/*γ* sequences from four CD8+ *γδ* TCRs (GD8-1, GD8-2, GD8-3, GD8-4) and two control TCRs (TB5, AT1G9) known to respond to HMB-PP. Jurkat *α*-*β*- cells expressing these TCRs were tested for reactivity to cell lines, Mtb-lysate, and HMB-PP, measured by the increase in CD69 expression 14 hours post stimulation. *P*-values were determined using one-way ANOVA followed by Dunnett’s multiple comparisons test. Error bars represent mean and standard deviations. **(e)** Percentage and number of CD8+ *γδ* T cells in unstimulated and Mtb-lysate (1*μ*g/ml)-stimulated tonsil organoid cultures (day7) from 3 healthy donors. **(f)** Clonal composition of CD8+ *γδ* T cells in unstimulated and Mtb-lysate-stimulated tonsil organoid cultures from one healthy donor (left). The total number of paired *γδ* TCRs is shown below each pie chart. For each TCR expressed by two or more cells (clonally expanded), the absolute number of cells expressing that TCR sequence is shown by a distinct colored section. The stacked bar plots show the V*γ*/*δ* chain pairings used by the clonally expanded tonsillar CD8+ *γδ* T cells in the unstimulated and Mtb-lysate stimulated cultures (right).

To determine the antigen specificity of the *γδ* T cell clones identified in controlled Mtb-infection, four clonally expanded TCRs with different *γδ* chain-pairings were selected for expression in Jurkat *α*-*β*- cells (**Fig. 4d**). Three of these clones responded to Mtb-lysate, but none of them to (E)-4-hydroxy-3-methyl-but-2-enyl pyrophosphate (HMBPP) (**Fig. 4d)**, a metabolite from the isoprenoid pathway in Mtb, which stimulates V*γ*9V*δ*2 T cells^31^. This observation indicated that *γδ* T cells induced at different stages of Mtb infection respond to different Mtb component(s).

It is plausible that these *γδ* T cells are induced at the site of infection and/or the draining lymph nodes. Given the difficulty in acquiring human lung and draining lymph node samples, to test this supposition, we utilized human tonsil organoids^32^ to evaluate Mtb-specific, lymphoid organ *γδ* T cell responses. We found that stimulation of tonsil organoid cultures with the Mtb-lysate resulted in increased frequencies and numbers of CD8+ *γδ* T cells in all 3 independent tonsil organoid cultures established from 3 healthy donors (**Fig. 4e**). Directly *ex vivo* single cell TCR sequencing of CD8+ *γδ* T cells isolated from a stimulated organoid culture demonstrated clonal expansion (**Fig. 4f**). ∼60% of the expanded clonotypes expressed V*δ*1 encoded TCRs. *γδ* T cells with other TCRs, including those encoded by V*γ*9V*δ*2 and with CDR3 sequences compatible for phosphoantigen recognition also showed clonal expansion in the tonsil organoid cultures (**Fig. 4f, Supplementary Table 4**). These observations are consistent with the idea that the Mtb-lysate consists of components which stimulate *γδ* T cells in both the acute and the controlled stages of infection, and suggest that Mtb-specific *γδ* T cell responses can be generated in lymphoid tissues regardless of the stage of infection.

Some *γδ* T cells induced in chronic CMV infection, as well as some Mtb-lysate responsive V*γ*9V*δ*2 T cells exhibit reactivity to tumor cell lines^15, 33^, such as T2 (lymphoblastoid cell line), K562 (leukemia cell line), and Daudi (lymphoma cell line), suggesting TCR reactivity to dysregulated host antigens. We therefore tested whether the TCRs expressed on *γδ* T cells induced in controlled Mtb infection have similar reactivities. We found that Clones GD8-1 and GD8-5 showed robust CD69 expression upon coculture with T2, but not K562 or Daudi cells. Clone GD8-3 was reactive to both T2 and Daudi, but not K562 cells, while clone GD8-2 recognized Daudi cells, but not T2 or K562 cells (**Fig. 4d**). These results indicate that while CD8+ *γδ* T cells as a group respond to antigens expressed by tumor cell lines, the TCRs show different fine specificities, which is expected from adaptive antigen recognition.

### γδ T cells that respond to controlled Mtb infection traverse distinct developmental trajectories

Given the observation that controlled and acute Mtb infection induced *γδ* T cells with different TCRs that respond to different components of Mtb-lysate, we sought to better understand the developmental/maturation paths that direct *γδ* T cells to the effector states in an infection stage-specific manner. To this end, we employed the newly developed Cytoskel algorithm^34^ to construct pseudotime trajectories which reflect cellular progression through specific activation and maturation pathways from single cell targeted mRNA expression and concomitant cell surface marker expression. Like many Trajectory-Inference (TI) methods, Cytoskel constructs a k-NN graph to capture local similarities between data cells. In contrast to other TI methods, Cytoskel constructs a minimum spanning tree (MST) connecting all the individual data cells as a subgraph of the k-NN graph and restricts trajectories to use only edges in the MST as these express the closest relations between cells. Branching trajectories are then constructed as extremal subtrees of the MST. This approach is different from the *γδ* T cell trajectory analysis described by Pizzolato et al., where V*δ*1+ and V*δ*2+ peripheral *γδ* T cells were isolated and analyzed separately and only linear pseudotemporal ordering of cells was considered^35^. To determine the developmental paths of *γδ* T cells with different TCRs in the controlled stage of Mtb infection, we performed concomitant single cell TCR determination, so that TCR usage could be associated with the trajectory analysis.

FACS sorted *γδ* T cells (N=24,888) from six donors with controlled Mtb infection were examined (**Extended Data Fig. 3 a, b**). We found that ∼10% of the cells had a naïve phenotype (defined as CD45RA+CD62L+CCR7+) (indicated as X on the map). These cells navigated a trajectory with a branch point (Branch Point 1) that led to two alternative developmental fates – trajectory 1 and trajectory 2, with similar numbers of cells in each trajectory. Trajectory 1 further bifurcated (Branch point 2) into two separate paths which ended at A and B, while trajectory 2 showed an early branching event (Branch Point 3) leading to two divergent cellular paths, eventually resulting in three branch termini (identified as C, D, and E). Additionally, some of the naïve *γδ* T cells appeared to travel a short distinct path (identified as F) that ran parallel to paths A and B and was considered a part of trajectory 1 (**Fig. 5a**). Notably, nearly all of the cells along trajectory 1 (comprised of Cell Clusters 2 and 3) appeared to be programmed to differentiate into cytolytic effectors with the expression of NK cell receptors, including CD16. While cells along paths A and B were CD8+, those along path F were CD8- (**Fig. 5b, Extended Data Fig. 4a**). In contrast, ∼85% of the cells in trajectory 2 appeared to be in a transitory state, trailing around Branch Point 3 (Cell Cluster 8) (**Fig. 5a**). Only ∼10% of the cells in trajectory 2 differentiated into cytolytic NK-like effectors, that also expressed CD16, but lacked CD8 expression (path C) (**Fig. 5b, Extended Data Fig. 4a**). The remaining ∼5% of cells followed the path from Branch Point 3 to terminus D and E via Branch Point 4 (Cell Clusters 7, 9, and 10). All three of these cell clusters expressed high levels of various cytokine receptors, including IL3RA, IL4R, IL23R, and IFNGR1 (**Extended Data Fig. 4b**). The end point E was very similar to D and appeared to be a side excursion of cells retracing their trails forward and backwards between these two paths. *γδ* T cells in Cell Cluster 7 expressed cytolytic effectors NKG7, GNLY, and the chemokine CCL5. *γδ* T cells in Cluster 9 developed into effectors that were distinguished by their expression of the cytokine IL-1*β* and chemokines, including CXCL2, CXCL3, CXCL5, and CXCL8 (IL-8), that target neutrophils and other myeloid cells (**Fig. 5c**). Clusters 7 and 9 also contained cells that expressed CD8 and CD16 (**Fig. 5b**).

**Figure 5.**
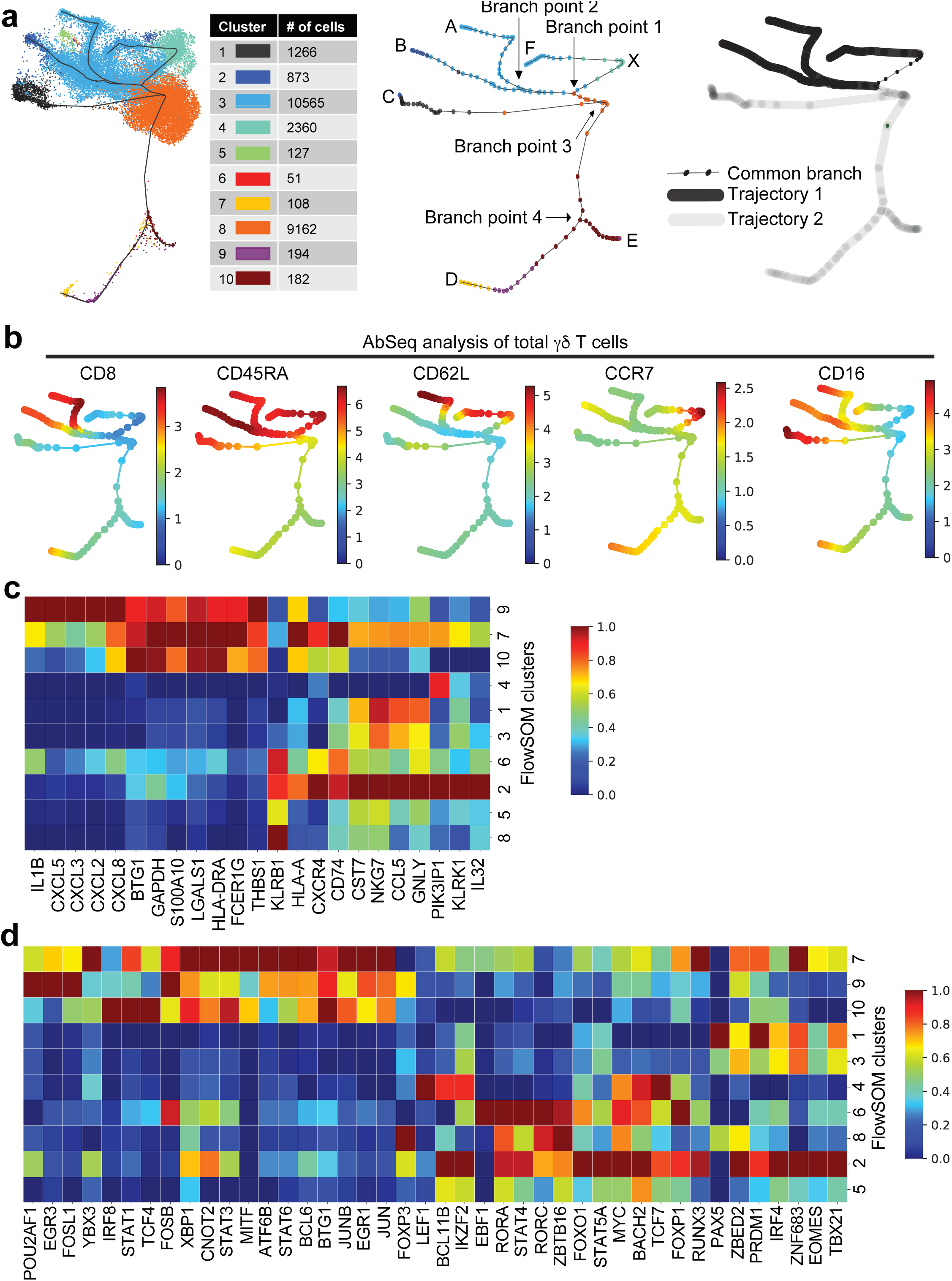
CD8+ γδ T cells traverse distinct differentiation trajectories in controlled Mtb infection. **(a)** Trajectory initiated UMAP plot of cell clusters with two-dimensional branching trajectories. Two-dimensional branching trajectories were constructed using the Cytoskel package (N=24,888 *γδ* T cells from 6 donors with controlled Mtb infection). The data cells were clustered into 10 clusters using the FlowSOM R package, applying the same distance calculation as the trajectory algorithm (left). The cells were colored based on FlowSOM clustering. Trajectory initialized UMAP starts with a 2D layout of trajectory cells using MDS. A modified UMAP algorithm is then run on the combined set of trajectory cells and data cells with the constraint that the trajectory cell positions remain fixed. Cluster 6 has very few cells (n=51) and is not visible since most of the cells are under other cells and scattered over a large area. Based on the expression of specific cell surface markers and genes, naïve cells were identified and marked as the starting point “X”, which bifurcated into two distinct trajectories. Each of the branch points are annotated and the branch termini are identified as A, B, C, D, E, and F (shown in the middle). The branches that comprise trajectory 1 are colored in black and those that constitute trajectory 2 are colored in gray (shown on the right). **(b)** Expression of selected cell surface markers along the trajectories as determined by antibody staining. The expression intensity of each marker is indicated, independently for each marker, by the colored gradient for which the range corresponds to the arcsinh transformed expression [arcsinh(x/5), where x=counts]. **(c, d)** Heatmaps of the expression level of ten most highly expressed genes (c) and transcription factors (d) in each cell cluster. For each cluster, the cluster mean was calculated for each gene, and then scaled across all clusters. The colored gradient thus represents the scaled expression of the mean expression of each gene measured across all clusters.

These divergent paths were marked by the expression of different transcription factors/related genes (**Fig. 5d**). The naïve point X, as expected, showed high expression of LEF1 and TCF7. In trajectory 1, the highly cytolytic endpoint B (Cell Cluster 2) was characterized by high levels of PRDM1 (BLIMP-1), TBX21 (T-bet), IRF4, and RUNX3 expression (**Fig. 5d**). A recent study has shown that RUNX3 synergizes with TCR signaling to establish IRF4-dependent transcriptional circuits that promote memory cytotoxic T lymphocyte formation^36^. These cells also expressed high levels of IKZF2 (Helios), BCL11B, FOXO1, and BACH2, transcription factors that function in promoting and stabilizing lineage commitment. In Trajectory 2, the branches leading to the end points D and E (Cell Clusters 7, 9, and 10), displayed enhanced AP-1 transcription factor associated FOSB, JUN, and JUNB expression, which are essential for the functional development of T cells, including IL-17-producing helper *αβ* T cells^37^. Consistent with the observation that some cells in Cluster 7 expressed cytolytic effector genes, they expressed many of the factors identified for cells in Cell Cluster 2, including PRDM1 (BLIMP-1), TBX21 (T-bet), EOMES, RUNX3, as well as ZNF683 (HOBIT), which is noted for promoting BLIMP-1 expression. Interestingly, *γδ* T cells in the transition state, Cluster 8, expressed (1) high levels of ZBTB16 (PLZF), which is commonly associated with *γδ* T cells that encounter ligands in the thymus, (2) RORA and RORC, factors associated with IL-17 making T cells and T regulatory cells, and (3) FOXP3, a lineage specification factor for T regulatory cells. The physiological relevance of this gene expression pattern will require further study. In addition, as our analysis was carried out with targeted scRNA-seq, it is conceivable that additional factors not listed in Fig. 5d, could be important for the cellular processes performed by these cells. Regardless, these data identify transcriptional regulators that may play instructive roles in *γδ* T cell effector fate decisions and maintenance.

Among the cells whose TCRs were successfully sequenced (N=12,706), 39%, 27%, and 26% comprised of V*δ*1+, V*γ*9-V*δ*2+, and V*γ*9+V*δ*2+ TCRs, respectively (**Extended data Fig. 5a**). The V*δ*1+ T cells made up 60% of cells in the naïve state (between X and Branch Point 1, Cell Cluster 4), and predominately (86% of the remaining V*δ*1+ cells) appeared in Trajectory 1. ∼90% of these V*δ*1+ cells traversed distinct differentiation paths (from Branch Point 2 to A and B, and X to F, Cell Cluster 3) leading to cytolytic effector cell fates. In sharp contrast, ∼73% of all V*γ*9+V*δ*2+ T cells (except those between X and Branch Point 1) belonged to trajectory 2, and ∼90% appeared to be in transitional states, gathered around branch points (between Branch Points 1 to 2, 1 to 3, and 3 to 4, Cell Cluster 8), with less than 10% progressing to differentiate into effectors at the various branch termini (C, D, and E). In this context, we found fetal-derived *γδ* T cells, defined as cells expressing TCRs without N-nucleotide additions in the CDR3 regions [annotated by IMGT/V-QUEST] in the PBMCs of all six donors (**Supplementary Table 5)**. ∼97% of these cells expressed V*γ*9+V*δ*2+ TCRs, frequently with the V*δ*2-D*δ*3-J*δ*3 rearrangement, and largely belonged to trajectory 2, located in Cell Cluster 8, a developmental stage distribution similar to other V*γ*9+V*δ*2+ cells (**Extended data Fig. 5b-e**). It is important to note that *γδ* T cells with different V*γ/*V*δ* TCRs showed preferential, but not exclusive distribution in any of the transition nodes and effector end points. This observation is consistent with the TCR usage described for cell clustering analysis from scRNA-seq data of *γδ* T cells from two adult PBMCs and two cord blood samples^38^.

Unlike the V*γ*9+V*δ*2+ T cells, the V*γ*9-V*δ*2+ T cells were equally abundant in trajectories 1 and 2, and both at branch points and branch termini. This is in line with reports that V*γ*9-V*δ*2+ and V*γ*9+V*δ*2+ T cells have different antigenic specificities and participate in different pathological conditions^11^. It also indicated that these cells would contribute to the cytolytic effector cell pool in controlled Mtb infection. Indeed, V*γ*9-V*δ*2+ cells comprised ∼30% of cells with cytolytic effector fate in trajectory 1 (from Branch Point 2 to A and B, and X to F), and approximately one third of cells with cytolytic effector fate in trajectory 2 (from Branch Point 3 to C).

In terms of clonal expansion of *γδ* T cells (based on *γδ* paired chain sequences) along the various branches, we found gradual increases in clonal expansion along trajectory 1 (Branch point 2 to A and B, and X to F) (**Extended Data Fig. 6, Supplementary Table 6**). In contrast, the cells in trajectory 2 showed limited clonal expansion, with no association between clonality and the pseudotemporal ordering of cells. In fact, BTG1 and LGALS1, two of the ten most abundantly expressed genes in Cell Clusters 7, 9, and 10 (**Fig. 5c**) are noted for their anti-proliferative function^39, 40^. These results were consistent with the TCR analysis of CD8+ and CD8- *γδ* T cells from the 4 donors described above (**Fig. 4a**).

Taken together, these results indicated that in the state of controlled Mtb infection, the responding *γδ* T cell population (predominantly the V*δ*1+ T cells) traversed distinct differentiation paths leading to cytolytic effector cell fates. By contrast, most of the V*γ*9V*δ*2 T cells were in transitory states, indicating that they are poised to develop into the highly cytolytic effectors reportedly found in acute Mtb infection^1^. These observations suggest that the cytolytic functional fate of *γδ* T cells in a given infection stage is acquired in the periphery as a consequence of antigen and environment driven events.

### Increased frequency of peripheral CD8+ γδ T cells is associated with chronic inflammatory conditions of diverse etiologies

The trajectory analysis indicated that *γδ* T cells traverse distinct effector differentiation paths to provide infection stage-specific response. To test whether increased circulating CD8+ *γδ* T cell frequencies may be a common feature of persistent or chronic infectious and inflammatory conditions, we analyzed flow cytometry data of PBMCs from cohorts of chronic HIV (adults), chronic cardiovascular disease (older adults) and acute influenza (adults) infection. We also analyzed publicly available datasets (https://flowrepository.org) from cohorts of melanoma (chronic)^41^ and COVID-19 (acute) patients. With striking consistency, circulating CD8+ *γδ* T cell frequency was found to increase across all cohorts presenting chronic or persistent inflammatory conditions, even the frequency of total *γδ* T cell remained unchanged (**Fig. 6a**). This trend was absent in donors with acute infections (**Fig. 6b**). Consistent with this finding, higher levels of circulating CD8+ *γδ* T cells have been reported for a small group of HIV seropositive subjects^42^, and in immunocompromised patients after allogeneic stem cell transplantation with CMV reactivation^15^.

**Figure 6.**
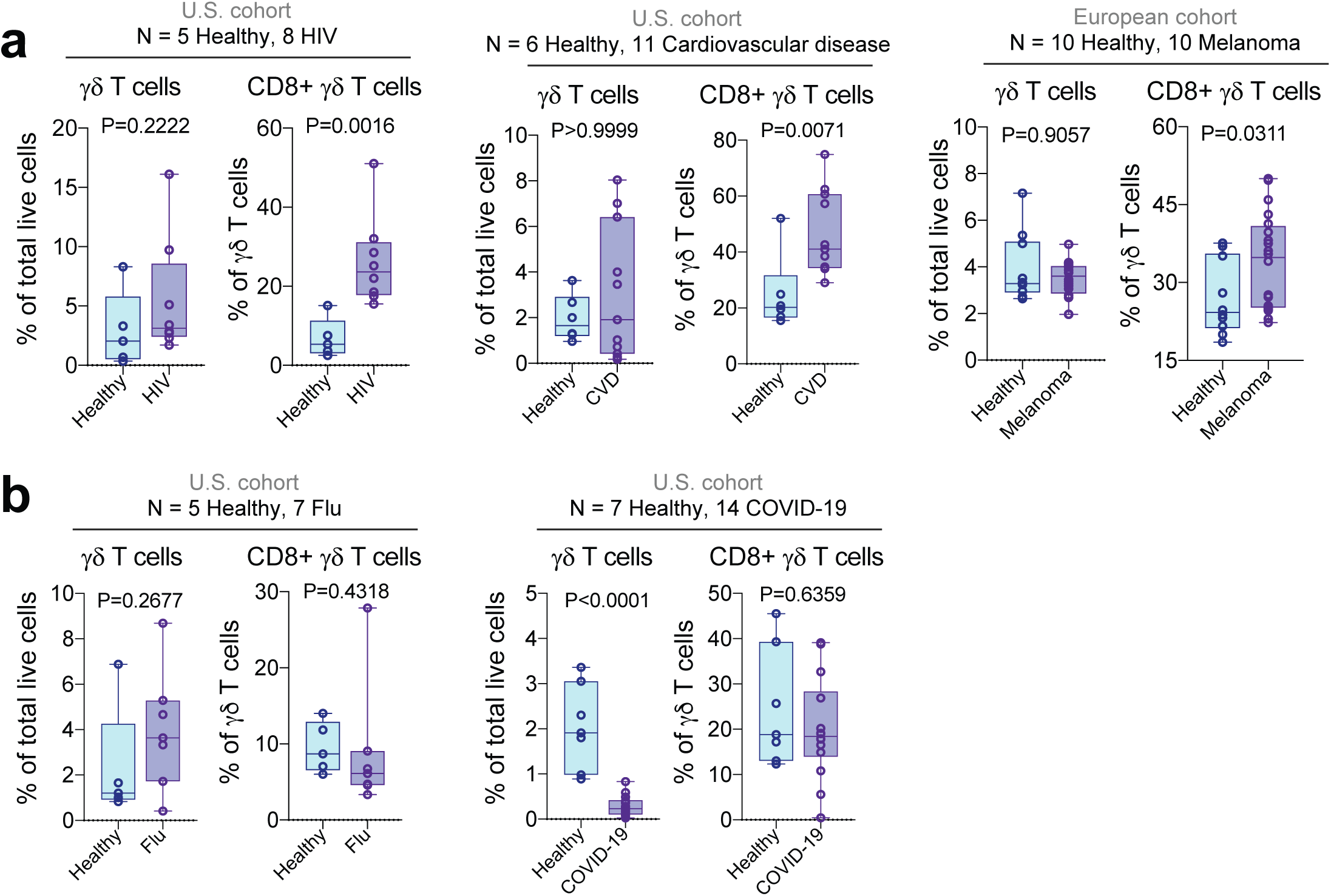
Increased frequency of circulating CD8+ γδ T cells associates with chronic inflammatory conditions of diverse etiologies. **(a, b)** Percentages of total *γδ* T cells and CD8+ *γδ* T cells in different cohorts of chronic (a) and acute (b) inflammatory conditions compared to healthy controls. *P*-values were determined using the Mann-Whitney test. Lower and upper hinges of boxes represent 25th to 75^th^ percentiles, the central line represents the median, and the whiskers extend to the highest and lowest values.

## Discussion

One difficulty of studying immune response in chronic Mtb infection is to identify cohorts that are truly in the designated infection state. Latent/persistent TB infection has been defined by having Mtb-specific *αβ* T cell IFN*γ* response in a blood sample, in the absence of clinical symptoms of active TB disease. Recent work has indicated that such responses may persist even after bacterial clearance^43^. It is possible that some of the individuals in our adolescent cohort have cleared the infection. However, these participants live in a setting with high endemic levels of Mtb and therefore have an increased likelihood of re-infections. In accordance with this, the results presented here show that most of the participants had circulating *γδ* T cells with phenotypes consistent with persistent antigen exposure, and most likely, persistent Mtb infection.

Our study highlights a key difference in how *αβ* T cells and *γδ* T cells contribute to host immune competence. In situations like chronic infections, cancer, and atherosclerosis, most antigen- specific effector *αβ* T cells in the presence of continuous TCR engagement progressively lose their effector functions and decrease in number, a phenomenon referred to as “exhaustion”. It is commonly considered that this diminished ability of *αβ* T cells to control the inflammatory response contributes to the establishment of chronic infection^44^. Here, we found that in controlled Mtb infection, an antigen driven response may expand *γδ* T cells and skew them towards terminally differentiated effectors. Although these cells have an attenuated response to TCR mediated signaling, they express NK cell receptors and have robust CD16-mediated ADCC response. This feature would allow antibodies, in addition to T cells, to contribute to the antigen specific response. In this context, it was reported that antibodies from individuals with persistent Mtb infection, have unique Fc functional profiles that promote selective binding to CD16, and can effectively drive intracellular Mtb killing^45^.

Indeed, development of CD16+ terminally differentiated cytolytic effector T cell response may be a general strategy for the host to control the infection. Chronic, untreated HIV-1 infection has been associated with elevated numbers of CD45RA+CD57+ terminal effector CD8 *αβ* T cells expressing CD16, and these CD16+ CD8 T cells mediate HIV-specific Ab-dependent cellular cytotoxicity activity at levels comparable with NK cells on a per cell basis^46^. We also found increased frequencies of cytolytic and polyfunctional CD16+ CD8+ *αβ* T cells in controlled Mtb infection in the same South African adolescent cohort studied here^16^.

In fact, in some chronic viral infections and cancer, CD8+ *αβ* T cells are found to traverse divergent trajectories – while some cells become dysfunctional and terminally exhausted^44^, others develop into effector memory cells^47^. Recent work has suggested that the different clonal trajectories are programmed by signaling through the TCRs^48–50^. Although it is unclear whether these effector *αβ* T cells express CD16, and can mount CD16-mediated response, these cells have characteristic features that are shared by the *γδ* T cells induced in controlled Mtb infection that we describe here. These include the lack of expression of phenotypic markers, CCR7, CD62L, CD28, CD27, and IL7R, sustained effector functions, and antigen driven increased cell numbers.

In fact, the NK-like functions adopted by the CD8+ *γδ* T cells in chronic or persistent infections may be driven by sustained antigenic stimulation. It has been reported that *in vitro* culture of PBMCs in the presence of IL-2 or IL-15 induces the expression of natural cytotoxicity receptors (NCRs) on V*δ*1+ *γδ* T cells. These *γδ* T cells express high levels of granzyme B and show cytotoxicity against leukemic and other types of neoplastic cells. Importantly, this process requires prolonged stimulation through the TCR and a functional phosphatidylinositol 3-kinase (PI- 3K)/AKT signaling pathway^51^. We found that in comparison to CD8- *γδ* T cells, CD8+ *γδ* T cells showed enhanced AKT phosphorylation and expressed higher levels of genes enriched in the (PI-3K)/AKT pathway and also expressed high levels of the IL-2R/IL-15R common *β* chain (**Extended Data Fig. 4b**). In addition, a population of *γδ* T cells with NK cell functions has also been described in mice^52^. These cells are characterized by very low cell surface TCR expression, but strong intracellular CD3 staining, suggesting ligand induced TCR internalization. These observations indicate the importance of antigen recognition in the induction of *γδ* T cell functionality, even when the response is triggered through receptors other than the TCR.

As mentioned above, our data shows that *γδ* T cells that expand at different stages of Mtb infection respond to different components of an Mtb-lysate, underscoring the importance of antigen specificity for *γδ* T cells to mount a context-dependent response. Yet, both sets of *γδ* T cells show reactivity to the same tumor cell lines. V*γ*9V*δ*2 T cells which are activated in acute/active Mtb respond to HMBPP, an intermediate of the methylerythritol phosphate (MEP) pathway of isoprenoid biosynthesis in pathogenic bacteria and parasite. V*γ*9V*δ*2 T cells also respond to IPP (isopentenyl pyrophosphate) and DMAPP (dimethylallyl pyrophosphate) derived from the mevalonate pathway of isoprenoid synthesis of eukaryotic cells. Both microbial and host derived phosphoantigens activate V*γ*9V*δ*2 T cells through the induction of a specific form of BTN2A1 and BTN3A1 expressed on the cell surface, which interact with the *γδ* TCRs^53, 54^. This ability to respond to metabolites generated in proliferating cells allows V*γ*9V*δ*2 T cells to be activated by tumor cells. Nonetheless, this is unlikely to be the molecular basis for the tumor cell line recognition by the CD8+ *γδ* T cells identified in controlled Mtb infection since they do not respond to HMBPP. Given the similarity between *γδ* TCRs and immunoglobulins in antigen recognition, and antigen specific repertoires^55^, it is possible that some of these *γδ* TCRs are cross-reactive, and like cross-reactive antibodies, they may respond to a broad range of antigens. In this respect, increased circulating CD8+ *γδ* T cells appears to be a common feature of persistent or chronic infectious and inflammatory conditions. Defining what triggers this subset of *γδ* T cells, not only in chronic infection but also in atherosclerosis and cancer will be an important task going forward.

## Author contributions

RRC and Y-hC conceptualized the study. RRC designed and performed experiments. JRV performed the Cytoskel analysis. OK performed bulk-RNA-seq data analysis. MO performed scTCR sequencing and Jurkat transductions. MS helped with scRNA-seq analysis. HH helped with the *in vitro* stimulation assays of PBMCs with Mtb-lysate. MD helped with mitochondrial mass assay. LvB, ES, XH, and PKN provided FACS data for HIV, Flu, and cardiovascular disease. TJS provided samples from the ACS cohort. MMD, TJS, and SB provided valuable insights. RRC and Y-hC conducted data analysis and wrote the manuscript with inputs from all authors.

## Acknowledgements

We thank the Stanford Human Immune Monitoring Core (HIMC) for conducting the scRNA-seq (BD Rhapsody experiments) and assisting with data analysis. This work was supported by the Bill and Melinda Gates Foundation (BMGF) (MMD, TJS, Y-hC), the Howard Hughes Medical Institute (MMD), the American Heart Association 181PA34170022, 20TPA35500081 (PKN), Global Health grants OPP1021972 and OPP1066265 (TJS), and the National Institutes of Health 5T32AI07290- 31 (RRC), K99 AI129739-02 (LvB), HL134830-01 (PKN), AI127128 (Y-hC). The Adolescent Cohort Study was study was also supported by Aeras and BMGF GC6-74 (grant 37772) and BMGF GC 12 (grant37885) for QuantiFERON testing.

## Methods

### Cohort descriptions

#### South African adolescent cohort study (ACS)

As previously described^16^, PBMCs from individuals who were uninfected or with controlled Mtb infection, aged 13-18 years, were used for CyTOF, flow-cytometry, scTCR-seq, bulk-RNA-seq, and targeted scRNA-seq analyses. All eligible donors themselves gave written informed assent while their parents/legal guardians gave written, informed consent. The study protocols were approved by the Human Research Ethics Committee of the University of Cape Town. Individuals were classified as Mtb-infected based on a positive QuantiFERON TB Gold In-tube assay (Qiagen; >0·35 IU/mL). All participants were healthy without signs or symptoms of active disease. Only adolescents who remained disease free for two years from the time of enrolment were included in the analysis.

#### U.S. adult cohorts

(a) Chronic cardiovascular disease: Peripheral blood samples were collected from older patients (aged above 55 years) who underwent heart transplantation, and age-matched healthy controls. All eligible subjects gave their informed and written consent. The study protocol was approved by the Stanford Institutional Review Board. (b) Influenza virus infection: The cohort were adult patients attending the emergency department or the express outpatient clinic at Stanford with flu-like symptoms. Influenza infection was confirmed by nasal swab. Healthy controls were adult blood donors from the Stanford Blood Bank. The study protocol was approved by the Stanford Institutional Review Board. (c) HIV infection: All HIV-1-infected adults provided written informed consent before participation in the study (www.clinicaltrials.gov;<u>NCT02018510</u>), which was conducted in accordance with Good Clinical Practice. Peripheral blood samples at baseline (i.e., before drug injection) were analyzed for this study. The protocol was approved by the Federal Drug Administration in the USA, the Paul Ehrlich Institute in Germany, and the Institutional Review Boards at the Rockefeller University and the University of Cologne. (d) Healthy tonsils for organoid cultures: Whole tonsils from individuals undergoing surgery for obstructive sleep apnea or hypertrophy were collected after written informed consent was obtained from adult participants. Ethics approval was granted by the Stanford University Institutional Review Board.

### CyTOF measurements and analysis

The antibody panels, staining protocols, and analysis methods used here have been thoroughly described in a previous study^16^. Briefly, PBMCs from the South African adolescent cohort were stained with two panels – one measuring 25 surface markers and 12 cytokines/effector molecules (cytokine panel) (N=14 uninfected and 14 controlled Mtb infection), and the other measuring 27 surface markers and 13 signaling effectors (phospho panel) (N=10 uninfected and 10 controlled Mtb infection). Data analysis was done on the Cytobank website (http://www.cytobank.org). First, total *γδ* T cells were manually gated for each sample and then Citrus analysis was performed to identify stratifying subsets between the infected and control groups. To determine differences in cell subset abundances, we used the SAM algorithm in Citrus, which assesses the false discovery rate (FDR) by permutations. We also used the dimensionality reduction technique – viSNE for the visualization of similarities and heterogeneity across individual cells. Manually gated *γδ* T cells from all samples were first concatenated and then visualized using viSNE. The analysis was performed on Cytobank.

### Flow cytometry measurements and analysis

For all flow cytometry experiments, PBMCs were blocked with Human TruStain FcX (Biolegend) for 20 minutes on ice and then stained on ice for 30 minutes with different antibody cocktails, all of which included anti-CD3 (UCHT1), anti-TCR*δ* (5A6.E9), anti-CD8 (RPA-T8), and anti-CD19 (HIB19) antibodies. Cells were washed with FACS buffer (PBS+ 1% FBS) and then analyzed on the LSRII or sorted using the BD Aria machines. Dead cells were defined as aqua-positive and were excluded from the analysis. All antibodies were validated by the manufacturers for flow cytometry application, as indicated on the manufacturer’s website. Data were analyzed using FlowJo version 10.2.

### Bulk-RNA-sequencing and analysis

CD8+ and CD8- *γδ* T cells from previously cryopreserved PBMC samples were directly FACS sorted into TRIZOL reagent from 5 donors with controlled Mtb infection (1000-5000 cells/sample) for RNA extraction. cDNA was generated using the SMART-Seq v4 Ultra Low Input RNA Kit (Takara). The amplified cDNA was fragmented, and libraries were made using the Illumina Nextera-XT kit, following manufacturer’s instructions. Raw Illumina sequences were checked for quality using FastQC version 0.11.8 (Babraham Bioinformatics) and aligned to the GRCh38/hg38 Human genome (UCSC Genome Browser) using star RNASeq aligner version 2.5.4b. Resulting alignments were processed with the feature Counts software version 2.0.0 (subread) to obtain raw counts for each gene. The raw counts were then analyzed using DESeq2 version 3.10 (Bioconductor) to get differential expression data as well as normalized counts.

### Mitochondrial mass detection

For the assessment of mitochondrial mass, we purchased the MitoTracker Green reagent from Invitrogen. The samples were stained according to manufacturer’s instructions. All cells were subsequently analyzed by flow cytometry.

### *In vitro* stimulation of PBMCs with anti-CD3 antibody and an Mtb-lysate

PBMCs were thawed in complete RPMI 1640 medium at 2×10^6^ cells per ml and recovered 12 hours before stimulation. PBMCs were stimulated with an Mtb-lysate (10 μg/ml) (obtained from BEI Resources) or plate-bound anti-CD3 antibody for 14 hours, and then stained with anti-CD3 (UCHT1), anti-TCR*δ* (5A6.E9), anti-CD8 (RPA-T8), anti-CD19 (HIB19), and anti-CD69 (FN50) antibodies. All antibodies were purchased from BioLegend. Dead cells were stained using the LIVE/DEAD Fixable Aqua Dead Cell Stain Kit (Thermo Fisher Scientific).

### FcγR-mediated antibody-dependent cell-mediated cytotoxicity (ADCC)

ADCC was carried out as previously described^56^, with some modifications. Briefly, P815 cells (a mouse leukemic cell line) were incubated with 10*μ*g/ml concentration of P815-specific monoclonal antibody, 2.4G2. Coated and uncoated P815 cells were then cocultured with FACS sorted and rested (24 hours) *γδ* T cells, or total PBMCs at an effector: target ratio of 1:10 in the presence of anti-CD107a (H4A3) antibody. After incubation, the cells were washed, stained, and CD107a degranulation was measured by cell acquisition on the LSRII machine.

### Direct *ex vivo* single cell γδ TCR determination

Barcode-enabled direct *ex vivo* single cell TCR determination was carried out and analyzed as described^28^. Briefly, the method involved single cell sorting of CD8+ and CD8- *γδ* T cells followed by reverse transcription and 3 progressive nested PCR reactions to amplify the CDR3 regions of both gamma and delta chains. The last PCR added barcodes to allow the identification of sequences derived from each individual cell studied. Amplified products from barcoded individual cells were combined and sequenced with the Illumina^TM^ MiSeq^TM^ platform. The resulting sequences were analyzed using VDJFasta, and the CDR3 nucleotide sequences were then extracted and translated. Human TCR sequencing primers are listed in **Supplementary Table 7**.

### Cell surface expression of γδ TCRs by lentiviral transduction and stimulation assays

Lentiviral transduction was performed as previously described^57^. Briefly, TCR *γ* and *δ* chain gene fragments were cloned into lentiviral constructs (nLV Dual Promoter EF-1a-MCS-PGK-Puro). For TCR expression, the TCR *γ* and *δ* chain constructs were transfected into 293X cells separately. The virus was collected after 72 hours of transfection and transduced into Jurkat *α*-/*β*- cells, which were selected for highest TCR expression by FACS sorting. 100 μl TCR transduced Jurkat *α*-/*β*- cells (10^6^ per ml) were co-cultured with 100 μl T2, K562, and Daudi cells (10^6^ per ml) in a 96-well plate. Different concentrations of plate-bound anti-CD3 stimulation was used as positive control. TCR transfectants in media only was used as negative control. HMB-PP and Mtb-lysate were used at 5*μ*M and 10*μ*g/ml concentrations, respectively. After 14-hour incubation, cells were collected and CD69 expression was measured using flow cytometry.

### Tonsil organoid culture experiments

Tonsil organoids were established as previously described^32^. Briefly, whole tonsils (overall healthy, without obvious signs of inflammation) were collected in saline after surgery and then immersed in an antimicrobial bath of Ham’s F12 medium (Gibco) containing Normocin (InvivoGen), penicillin and streptomycin for 1 hour at 4 °C for decontamination of the tissue. Tonsils were then briefly rinsed with PBS and manually disrupted into a suspension by processing through a 100-μm strainer with a syringe plunger and cryopreserved. Frozen cells were thawed, washed, enumerated, and then plated (6X10^6^ cells in 100 μl per well) into permeable (0.4-μm pore size) membranes placed in standard 12-well tissue-culture plates. Mtb-lysate (1μg/ml) was then directly added to the cells and cultured for 7 days. Organoids were harvested 7 days post stimulation, cells were washed and then used for flow cytometry analysis or single-cell sorting for TCR determination.

### BD Rhapsody single-cell analysis

*γδ* T cells were FACS-sorted from six donors with controlled Mtb infection. Samples were stained with oligonucleotide-conjugated Sample Tags from the BD Human Single-Cell Multiplexing Kit in BD staining buffer following the manufacturer’s protocol. Barcoded samples were then washed and spun down at 350xg for 10 minutes and pooled. Pooled sample was then stained concurrently with a panel of 12 oligonucleotide-conjugated antibodies: anti-CD3 (SK7), anti-CD8 (RPA-T8), anti-CD16 (3G8), anti-CCR7 (2-L1-A), anti-CD62L (DREG-56), anti-CD45RA (HI100), anti-CD45RO (UCHL1), anti-CD158e1/KIR-NKB1 (DX9), anti-CD335 (9E2), anti-CD337/NKp30 (P30-15), anti-KIR-NKAT2 (DX27), and anti-NKp44 (P44-8) from BD. Staining was performed using the BD staining buffer for 30 minutes on ice, samples were then spun down at 350xg for 10 minutes and then washed three times. Pellet was resuspended in Rhapsody buffer for sort and capture. Cell capture and library preparation were completed using the BD Rhapsody Targeted mRNA and AbSeq Reagent kits. Briefly, cells were captured with beads in a microwell plate, followed by cell lysis, bead retrieval, cDNA synthesis, template switching, Klenow extension, and library preparation following the BD Rhapsody protocol in the Stanford Human Immune Monitoring Center. Libraries were prepared for T cell receptor, sample tags, targeted mRNA using the BD standard Immune Response panel, and AbSeq. Sequencing was completed on NovaSeq (Illumina, San Diego, CA) in the Stanford Genome Sequencing Service Center, and at Novogene (US Davis, CA).

Data were processed using the Seven Bridges Genomics online platform (San Francisco, CA) and BD Rhapsody Targeted Analysis Pipeline with V(D)J processing incorporated. After processing, data were imported into SeqGeq version 1.6.0 (BD, Ashland, OR). The import included a CSV file of all the data, and CSV files identifying the Sample Tag and V(D)J calls. Then the plug-in Lex BDSMK was run to separate out the Sample Tags, and the VDJ Explorer to identify clones. Data was further processed in Seurat pipeline^58^ to annotate cell subsets by clustering algorithm, and remove contaminating B cells, alpha beta T cells, and some myeloid cells. The data was further cleaned of various outlier cells – these consisted of cells with CD22 mRNA counts greater than 1 (6 cells), and cells which had a gene with an mRNA count greater than 500, and were identified as outliers (204 cells). The final data set consisted of 24,888 cells with each cell having counts for 12 proteins and 399 transcripts.

### Pseudotime Trajectories analysis

The data from Rhapsody single-cell analysis was transformed by replacing each count x by arcsinh(x/5). We refer to the D = 411 transformed coordinates (12 cell surface markers and 399 transcripts) for each cell as the feature coordinates and the D dimensional space as feature space. The data cells form a cloud of points in the feature space. Branching trajectories were then constructed using the cytoskel package. Briefly cytoskel constructs a k-nearest neighbor graph (k-NN graph) with edges connecting cells labeled by distances -dissimilarities between data points. This step captures the local and global structure of the cell cloud. Distances are simply the Euclidean distances between cells in the feature space. For the current data set we used k = 30 neighbors. Cytoskel then constructs a minimum spanning tree (MST) of the k-NN graph. The MST is the set of cell connections with the shortest total length which connect all cells. In the MST, each cell is only connected to the minimum set of cells with greatest similarity to itself such that all cells are connected.

Branch or trajectory construction then proceeds by first finding the two cells which are furthest apart as measured along the edges of the MST and constructing the graph path joining them. The next path segment is set of cells and edges from this first existing path to the cell furthest from the existing path. The process is then repeated a specified number of times. The result is a trajectory tree graph linking some subset of the data cells. An averaging step is carried out. The data cells are duplicated, and the coordinates of each duplicated cell are replaced by an average over near neighbors out to some distance in the MST. The averaged cells are referred to as pseudo-cells. The pseudo-cells which are part of the found trajectories are also referred to as trajectory cells. In the case of scRNA-seq data, this averaging is similar to imputation as done in MAGIC^59^ and kNN smoothing^60^.

The trajectories are represented as linked pseudo-cells. No assumption is made in the algorithm about the starting point of the trajectories. After trajectory construction the user can specify the starting branch. Given the starting branch, pseudo-time can be assigned to the trajectory pseudo-cells. Original data cells are associated with their closest pseudo-cell. Two dimensional plots of the pseudo-cell trajectories were constructed using metric multidimensional scaling (MDS) based on distances between trajectory cells in the original N dimensional feature space. This method creates a two-dimensional layout of the trajectory data points while trying to preserve as closely as possible the N dimensional distances between the data points. The cells in the plot are then colored by feature values of interest. This method clearly shows relationships between the trajectory branches and progression of features along the trajectories.

### FlowSOM Clustering

We performed FlowSOM^61^ meta-clustering on the data cells. FlowSOM first constructs a self-organizing map (SOM) which is a set of cell groups arranged on a (for example) 2D 10 by 10 grid such that the cells in each group are similar to each other and nearby groups are more similar than distant groups. The groups are then joined by a minimum spanning tree between groups. Finally, higher level clustering is performed on the groups forming meta-clusters each of which contains one or more low level groups. In the following we refer to the meta-clusters simply as clusters. The data cells were clustered into 10 meta-clusters using the FlowSOM R package. The clustering used the same distance calculation as the trajectory algorithm. For each cluster, the cluster mean was calculated for each gene. For the subset of markers of interest, a dataframe was constructed with each row containing the average values for each marker of interest for a given cluster. This dataframe was passed to the clustermap function of the Python seaborn plotting package with scaling set so that the maximum value in each column was scaled to a value of 1.0 to make the gene expression differences and similarities between the clusters clearer.

---

**Extended Data Figure 1.**
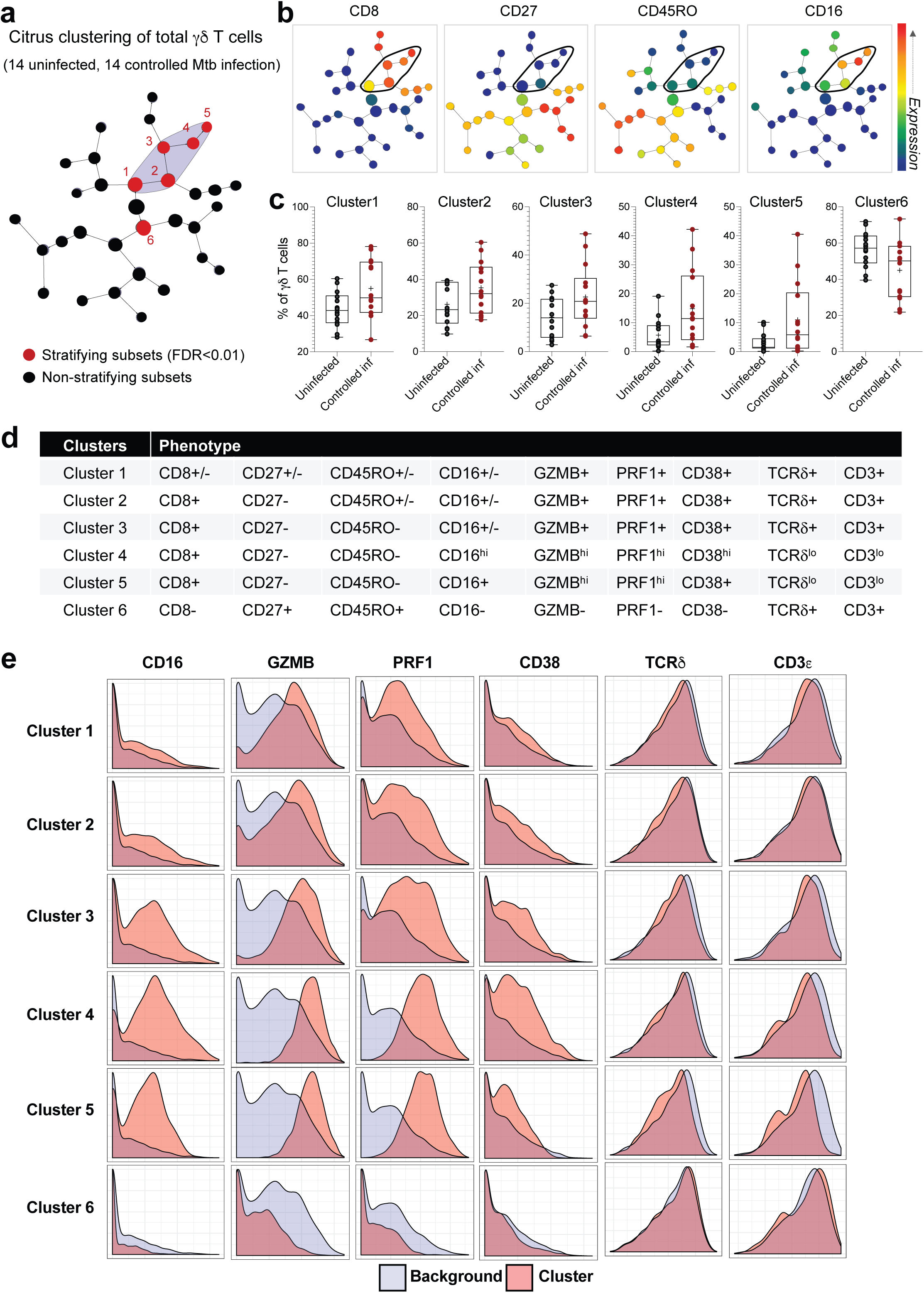
Increased frequency of peripheral CD8+ γδ T cells at controlled stage of Mtb infection. **(a)** Peripheral *γδ* T cells from 14 individuals with controlled Mtb infection and 14 uninfected participants of a South African adolescent cohort were examined for their surface and cytokine/effector molecule expression and analyzed by an unsupervised hierarchical clustering approach (Citrus). Cell clusters are represented as nodes (circles) in this Citrus-derived circular dendrogram, which delineates lineage relationships that were identified from the data. Cluster granularity (that is, cell subset specificity) increases from the center of the diagram to the periphery. Citrus plots showing, based on protein expression, clusters (in red, designated 1-5) that exhibit significantly different abundances (SAM analysis, FDR<1%) between the uninfected donors and individuals with controlled Mtb infection. **(b)** Annotation of cluster hierarchy plots based on surface marker expression. The expression intensity of each marker used for cell population characterization is overlaid per cluster on the Citrus circular dendrogram and is indicated, independently for each marker, by the colored gradient for which the range corresponds to the arcsinh transformed expression of the median marker expression measured across all Citrus clusters. **(c)** Percentages of the stratifying clusters in uninfected and donors with controlled Mtb infection. **(d)** Phenotype of the *γδ* T cells in each of the stratifying clusters. **(e)** Histograms showing the expression of selected markers on individual Citrus clusters (in orange) relative to the background (in grey).

**Extended Data Figure 2.**
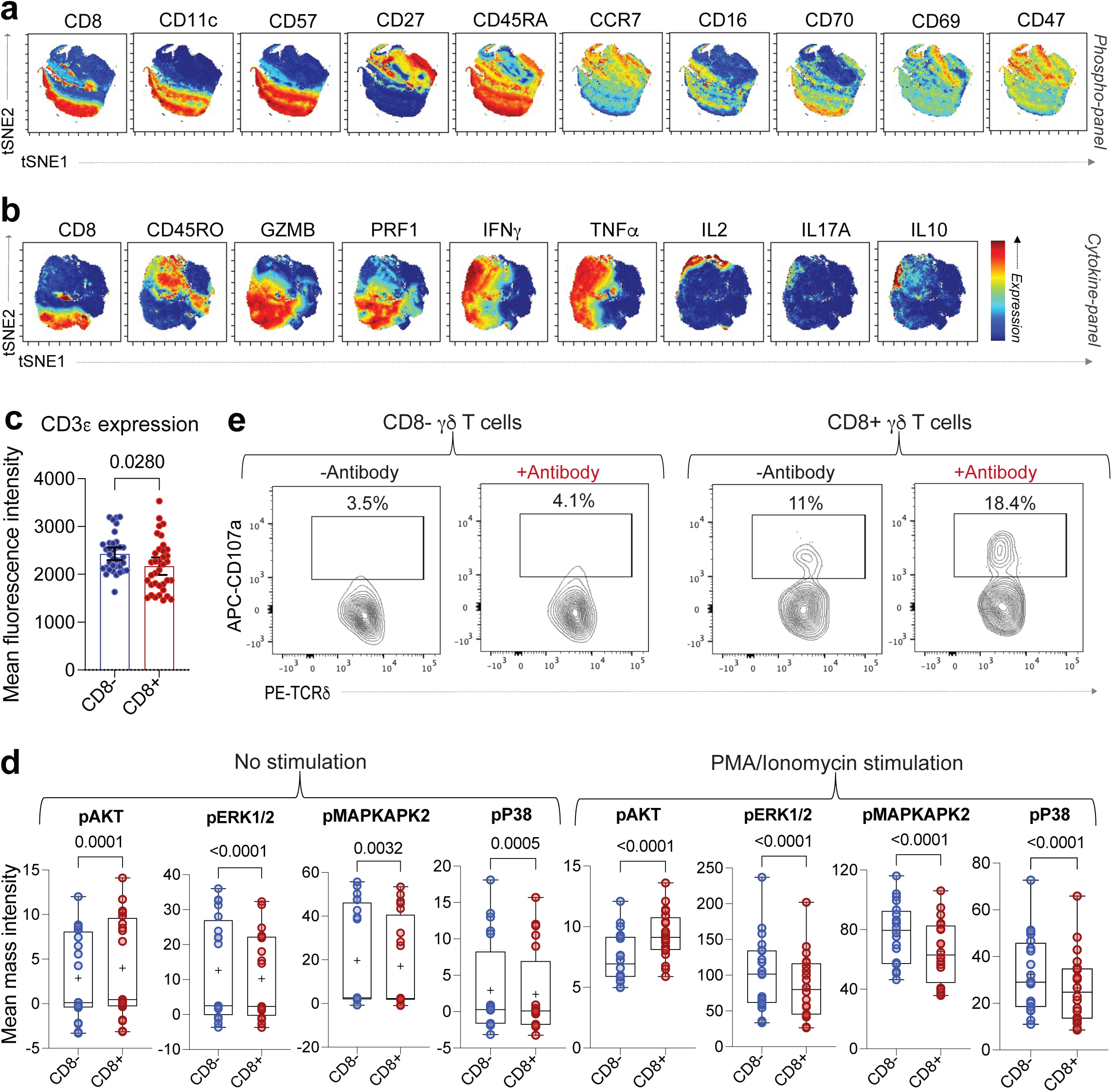
Peripheral CD8+ γδ T cells are terminally differentiated cytolytic effectors with characteristics of persistent antigenic exposure, distinct signaling potential and mount more robust ADCC response than CD8-γδ T cells. **(a, b)** viSNE analysis of mass cytometry data showing the protein expression of selected surface markers on peripheral *γδ* T cells from the phospho-panel (a) and cytokine-panel (b). **(c)** Mean fluorescence intensity of cell surface CD3*ε* expression on CD8- and CD8+ *γδ* T cells measured by flow cytometry analysis of 17 uninfected donors and 19 donors with controlled Mtb infection. *P*-value was derived using Wilcoxon matched-pairs signed rank test. Error bars represent mean and 95% confidence intervals. **(d)** Mean mass intensities of selected phospho-signaling effectors in CD8- and CD8+ *γδ* T cell subsets (N=20 samples/subset) at baseline (unstimulated condition) and post PMA- ionomycin stimulation. *P*-values were derived using Wilcoxon matched-pairs signed rank test. Lower and upper hinges of boxes represent 25th to 75^th^ percentiles, the central line represents the median, the plus sign represents the mean, and the whiskers extend to the highest and lowest values. **(e)** Contour plots showing the ADCC potential of CD8- and CD8+ *γδ* T cells, measured by the expression of CD107a in the presence of antibody-coated or uncoated P815 target cells.

**Extended Data Figure 3.**
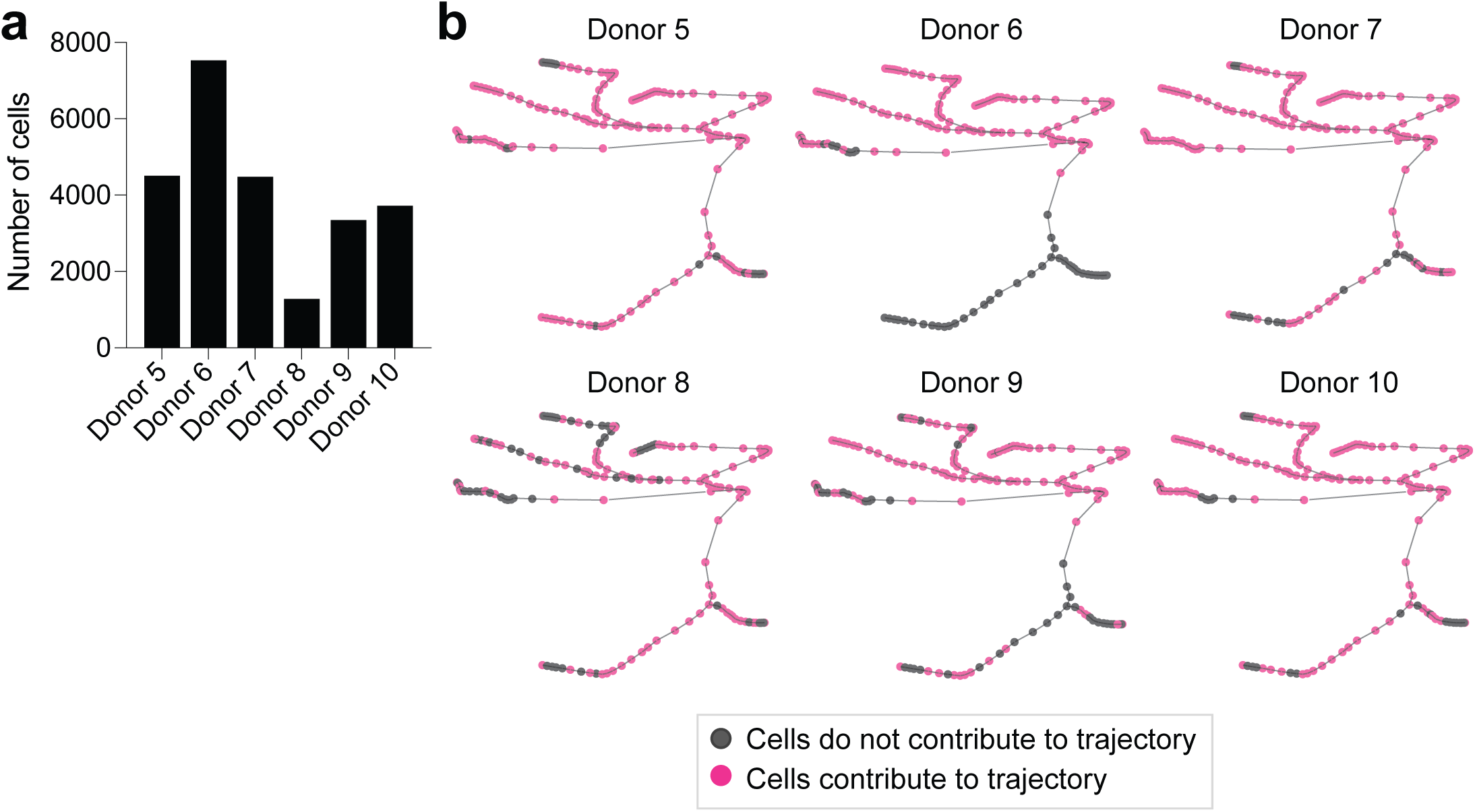
The establishment of peripheral γδ T cells trajectory analysis based on BD Rhapsody data. **(a)** Number of cells from each of the six donors with controlled Mtb infection that contributed to the Cytoskel trajectory analysis. **(b)** Two-dimensional Cytoskel plots showing that cells from each donor contribute to the construction of the various branches in the trajectory analysis. Pink represents parts of the trajectory that include cells from a given donor, while gray represents areas that do not include cells from a given sample.

**Extended Data Figure 4.**
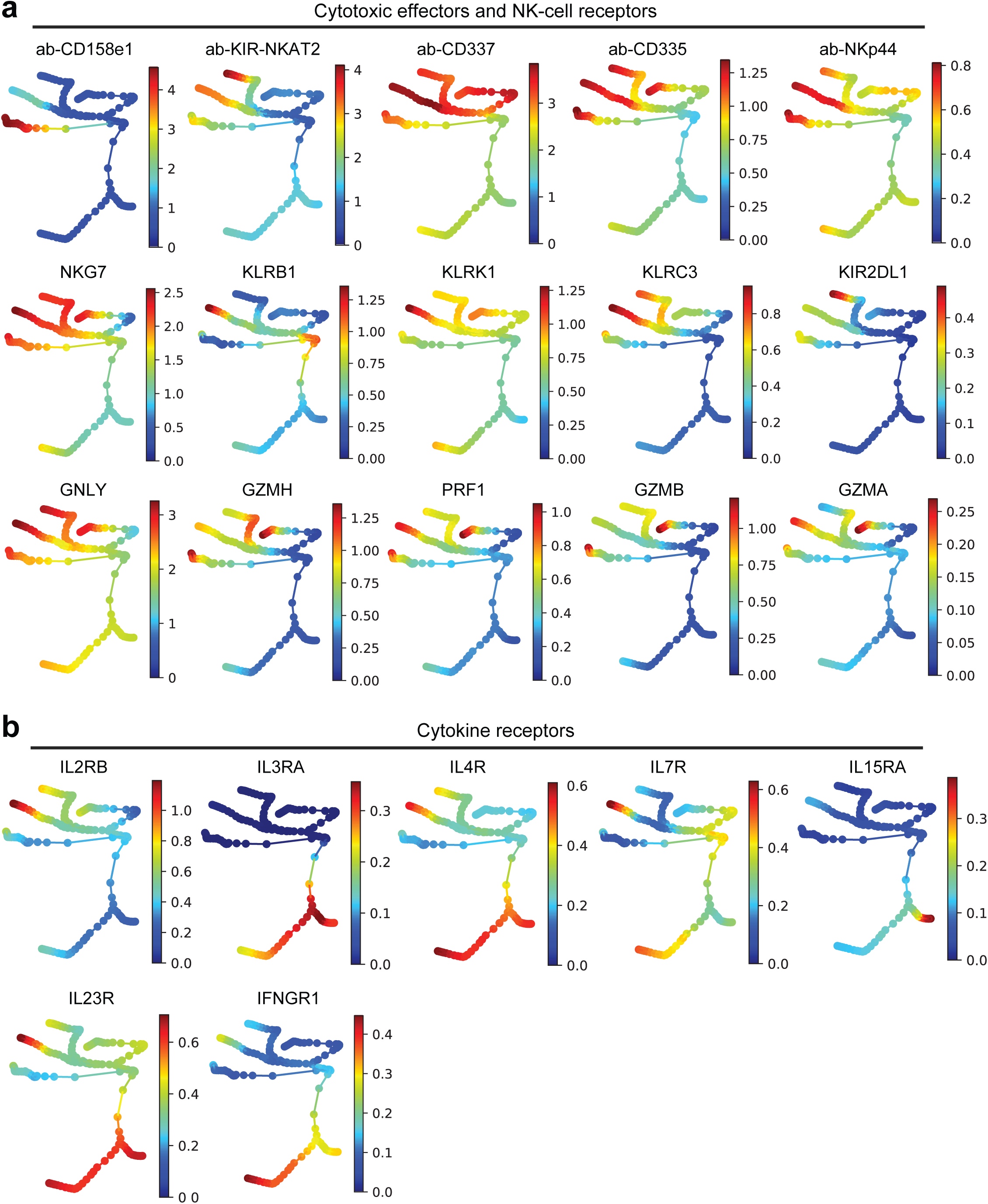
Pseudotemporal reconstruction of peripheral γδ T cells shows changes in gene expression patterns along the various differentiation trajectories. **(a-c)** Two-dimensional Cytoskel trajectory plots showing the normalized gene expression of selected cytolytic effectors and NK-cell receptors (a) and cytokine receptors (b) along the different *γδ* T cell effector differentiation branches. Normalized expression represents arcsinh(x/5), where x=counts.

**Extended Data Figure 5.**
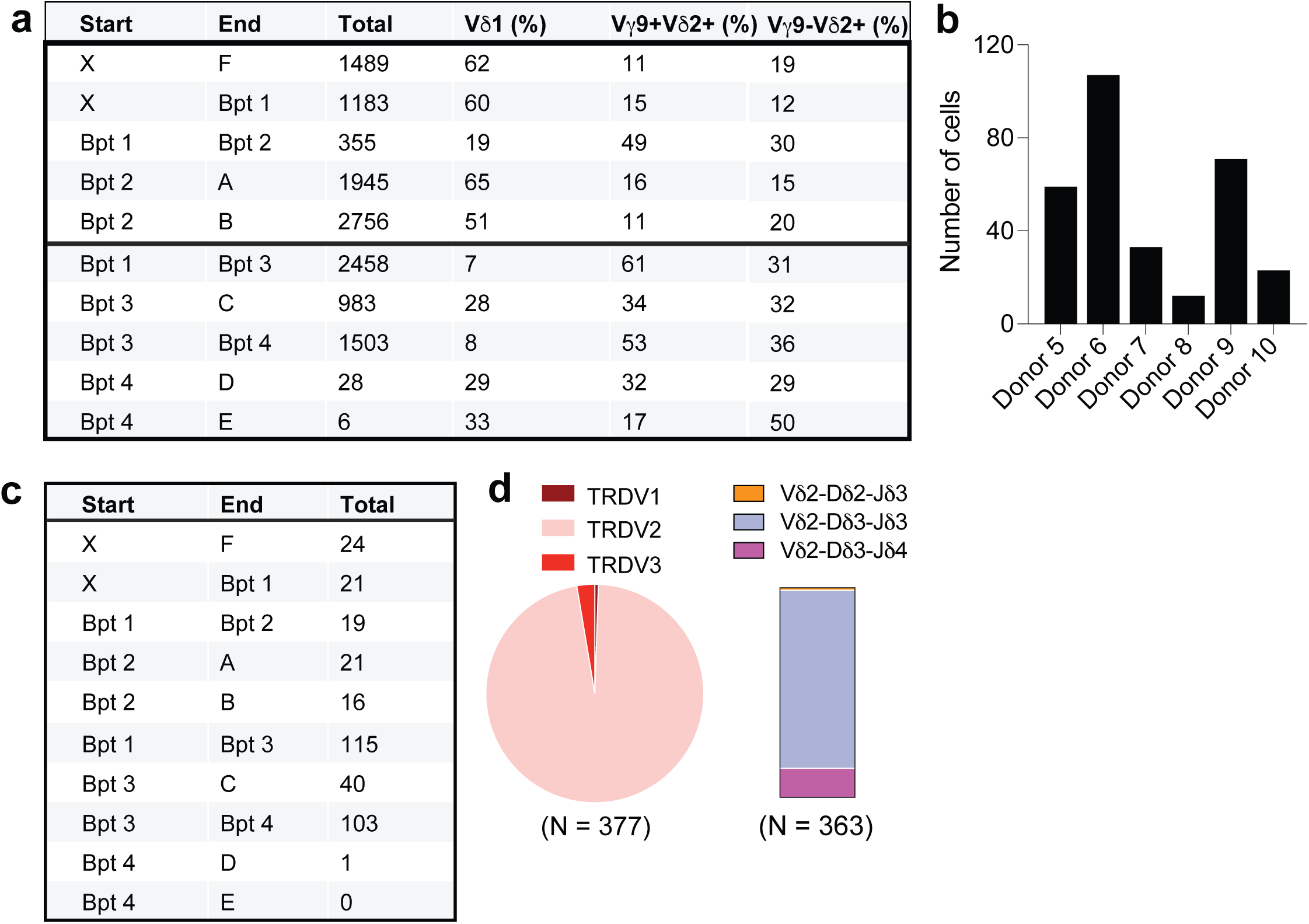
Distribution of γδ T cells with different TCRs along the trajectories. **(a)** Number of V*δ*1+, V*γ*9+V*δ*2+, and V*γ*9-V*δ*2+TCR sequences found along the trajectories. **(b)** Number of fetal-derived sequences found among peripheral *γδ* T cells in each of the six donors with controlled Mtb infection. CDR3*δ* sequences were analyzed in IMGT/V-QUEST and classified as “fetal-derived” if no N-nucleotides were present. **(c)** Number of fetal-derived *γδ* TCR sequences found along the different Cytoskel trajectories. **(d)** V*δ*-gene usage by all fetal-derived *γδ* T cells (N=377) (left), and D*δ−*J*δ* usage by V*δ*2-expressing fetal-derived *γδ* T cells (N=363) (right).

**Extended Data Figure 6.**
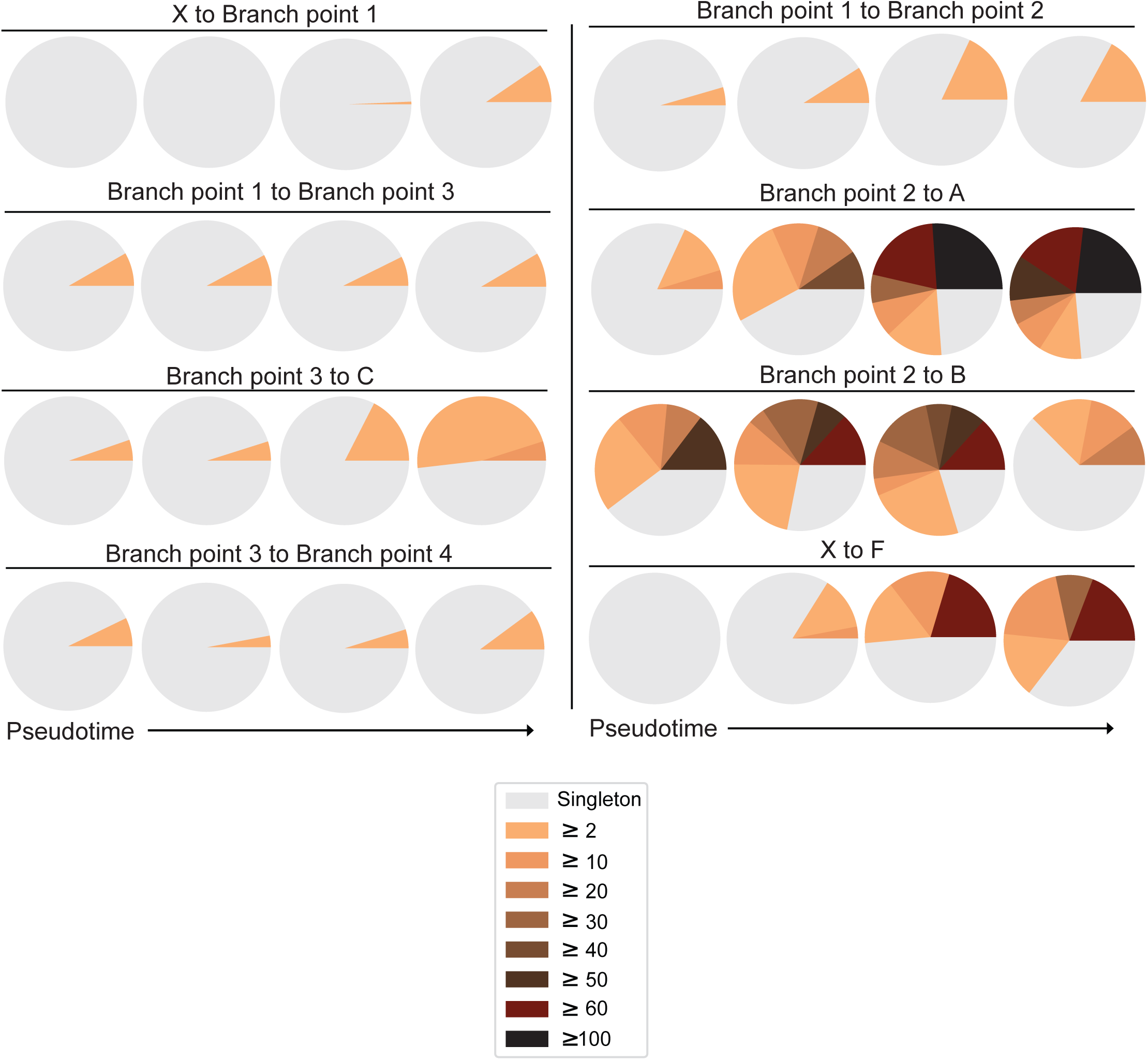
Clonal expansion patterns along γδ T cell differentiation trajectories. Pie charts showing the clonal expansion of *γδ* T cells along pseudotime axes. Cells along each Cytoskel branch were grouped into 4 pseudotime bins of approximately equal number of TCR sequences. For each TCR clone expressed by two or more cells (clonally expanded), the absolute number of cells expressing that clone is shown by a distinct colored section.

## References

1 Chen, Z. W. & Letvin, N. L. Vγ2Vδ2+ T cells and anti-microbial immune responses. Microbes and Infection 5, 491–498, doi:10.1016/s1286-4579(03)00074-1 (2003).

2 Barcy, S. et al. Gamma delta+ T cells involvement in viral immune control of chronic human herpesvirus 8 infection. J Immunol 180, 3417–3425, doi:10.4049/jimmunol.180.5.3417 (2008).

3 Spencer, C. T. et al. Granzyme A produced by gamma(9)delta(2) T cells induces human macrophages to inhibit growth of an intracellular pathogen. PLoS Pathog 9, e1003119, doi:10.1371/journal.ppat.1003119 (2013).

4 Junqueira, C. et al. gammadelta T cells suppress Plasmodium falciparum blood-stage infection by direct killing and phagocytosis. Nat Immunol 22, 347–357, doi:10.1038/s41590-020-00847-4 (2021).

5 Couzi, L. et al. Antibody-dependent anti-cytomegalovirus activity of human gammadelta T cells expressing CD16 (FcgammaRIIIa). Blood 119, 1418–1427, doi:10.1182/blood-2011-06-363655 (2012).

6 Caccamo, N. et al. Differentiation, phenotype, and function of interleukin-17-producing human Vgamma9Vdelta2 T cells. Blood 118, 129–138, doi:10.1182/blood-2011-01-331298 (2011).

7 Peng, M. Y. et al. Interleukin 17-producing gamma delta T cells increased in patients with active pulmonary tuberculosis. Cell Mol Immunol 5, 203–208, doi:10.1038/cmi.2008.25 (2008).

8 Papotto, P. H., Ribot, J. C. & Silva-Santos, B. IL-17(+) gammadelta T cells as kick-starters of inflammation. Nat Immunol 18, 604–611, doi:10.1038/ni.3726 (2017).

9 Khairallah, C., Dechanet-Merville, J. & Capone, M. gammadelta T Cell-Mediated Immunity to Cytomegalovirus Infection. Front Immunol 8, 105, doi:10.3389/fimmu.2017.00105 (2017).

10 Kaminski, H. et al. Understanding human gammadelta T cell biology toward a better management of cytomegalovirus infection. Immunol Rev 298, 264–288, doi:10.1111/imr.12922 (2020).

11 Kaminski, H. et al. Characterization of a Unique gammadelta T-Cell Subset as a Specific Marker of Cytomegalovirus Infection Severity. J Infect Dis 223, 655–666, doi:10.1093/infdis/jiaa400 (2021).

12 Pitard, V. et al. Long-term expansion of effector/memory Vdelta2-gammadelta T cells is a specific blood signature of CMV infection. Blood 112, 1317–1324, doi:10.1182/blood-2008-01-136713 (2008).

13 De Maria, A. et al. Selective increase of a subset of T cell receptor gamma delta T lymphocytes in the peripheral blood of patients with human immunodeficiency virus type 1 infection. J Infect Dis 165, 917–919, doi:10.1093/infdis/165.5.917 (1992).

14 Farnault, L. et al. Clinical evidence implicating gamma-delta T cells in EBV control following cord blood transplantation. Bone Marrow Transplant 48, 1478–1479, doi:10.1038/bmt.2013.75 (2013).

15 Scheper, W. et al. gammadeltaT cells elicited by CMV reactivation after allo-SCT cross-recognize CMV and leukemia. Leukemia 27, 1328–1338, doi:10.1038/leu.2012.374 (2013).

16 Roy Chowdhury, R. et al. A multi-cohort study of the immune factors associated with M. tuberculosis infection outcomes. Nature 560, 644–648, doi:10.1038/s41586-018-0439-x (2018).

17 Mahomed, H. et al. Predictive factors for latent tuberculosis infection among adolescents in a high-burden area in South Africa. International Journal of Tuberculosis and Lung Disease 15, 331–336 (2011).

18 Bruggner, R. V., Bodenmiller, B., Dill, D. L., Tibshirani, R. J. & Nolan, G. P. Automated identification of stratifying signatures in cellular subpopulations. Proc Natl Acad Sci U S A 111, E2770–2777, doi:10.1073/pnas.1408792111 (2014).

19 Huleatt, J. W. & Lefrancois, L. Antigen-driven induction of CD11c on intestinal intraepithelial lymphocytes and CD8+ T cells in vivo. J Immunol 154, 5684–5693 (1995).

20 Brugnoni, D. et al. CD70 expression on T-cell subpopulations: study of normal individuals and patients with chronic immune activation. Immunology Letters 55, 99–104, doi:10.1016/s0165-2478(96)02693-4 (1997).

21 Macintyre, A. N. et al. Protein kinase B controls transcriptional programs that direct cytotoxic T cell fate but is dispensable for T cell metabolism. Immunity 34, 224–236, doi:10.1016/j.immuni.2011.01.012 (2011).

22 Kersh, E. N. et al. TCR signal transduction in antigen-specific memory CD8 T cells. J Immunol 170, 5455–5463, doi:10.4049/jimmunol.170.11.5455 (2003).

23 Khawar, B., Abbasi, M. H. & Sheikh, N. A panoramic spectrum of complex interplay between the immune system and IL-32 during pathogenesis of various systemic infections and inflammation. Eur J Med Res 20, 7, doi:10.1186/s40001-015-0083-y (2015).

24 Ordway, D. et al. XCL1 (lymphotactin) chemokine produced by activated CD8 T cells during the chronic stage of infection with Mycobacterium tuberculosis negatively affects production of IFN-gamma by CD4 T cells and participates in granuloma stability. J Leukoc Biol 82, 1221–1229, doi:10.1189/jlb.0607426 (2007).

25 Butcher, E. C. & Picker, L. J. Lymphocyte homing and homeostasis. Science 272, 60–66, doi:10.1126/science.272.5258.60 (1996).

26 Katagiri, K. & Kinashi, T. Rap1 and integrin inside-out signaling. Methods Mol Biol 757, 279–296, doi:10.1007/978-1-61779-166-6_18 (2012).

27 Geltink, R. I. K., Kyle, R. L. & Pearce, E. L. Unraveling the Complex Interplay Between T Cell Metabolism and Function. Annu Rev Immunol 36, 461–488, doi:10.1146/annurev-immunol-042617-053019 (2018).

28 Han, A. et al. Dietary gluten triggers concomitant activation of CD4+ and CD8+ alphabeta T cells and gammadelta T cells in celiac disease. Proc Natl Acad Sci U S A 110, 13073–13078, doi:10.1073/pnas.1311861110 (2013).

29 Cheng, C. et al. Next generation sequencing reveals changes of the gammadelta T cell receptor repertoires in patients with pulmonary tuberculosis. Sci Rep 8, 3956, doi:10.1038/s41598-018-22061-x (2018).

30 Dechanet, J. et al. Implication of gammadelta T cells in the human immune response to cytomegalovirus. J Clin Invest 103, 1437–1449, doi:10.1172/JCI5409 (1999).

31 Morita, C. T., Jin, C., Sarikonda, G. & Wang, H. Nonpeptide antigens, presentation mechanisms, and immunological memory of human Vgamma2Vdelta2 T cells: discriminating friend from foe through the recognition of prenyl pyrophosphate antigens. Immunol Rev 215, 59–76, doi:10.1111/j.1600-065X.2006.00479.x (2007).

32 Wagar, L. E. et al. Modeling human adaptive immune responses with tonsil organoids. Nat Med 27, 125–135, doi:10.1038/s41591-020-01145-0 (2021).

33 Bukowski, J. F. et al. V gamma 2V delta 2 TCR-dependent recognition of non-peptide antigens and Daudi cells analyzed by TCR gene transfer. J Immunol 154, 998–1006 (1995).

34 John R. Valainis, S. C. K., Sean C. Bendall. Cytoskel: trajectory inference and analysis. dx.doi.org [10.5281/zenodo.4818819], doi:10.5281/zenodo.4818819 (2021).

35 Pizzolato, G. et al. Single-cell RNA sequencing unveils the shared and the distinct cytotoxic hallmarks of human TCRVdelta1 and TCRVdelta2 gammadelta T lymphocytes. Proc Natl Acad Sci U S A 116, 11906–11915, doi:10.1073/pnas.1818488116 (2019).

36 Wang, D. et al. The Transcription Factor Runx3 Establishes Chromatin Accessibility of cis-Regulatory Landscapes that Drive Memory Cytotoxic T Lymphocyte Formation. Immunity 48, 659–674 e656, doi:10.1016/j.immuni.2018.03.028 (2018).

37 Yamazaki, S. et al. The AP-1 transcription factor JunB is required for Th17 cell differentiation. Sci Rep 7, 17402, doi:10.1038/s41598-017-17597-3 (2017).

38 Tan, L. et al. A fetal wave of human type 3 effector gammadelta cells with restricted TCR diversity persists into adulthood. Sci Immunol 6, doi:10.1126/sciimmunol.abf0125 (2021).

39 Rouault, J. P. et al. BTG1, a member of a new family of antiproliferative genes. EMBO J 11, 1663–1670 (1992).

40 Blaser, C. et al. β-Galactoside-binding protein secreted by activated T cells inhibits antigen-induced proliferation of T cells. European Journal of Immunology 28, 2311–2319, doi:10.1002/(sici)1521-4141(199808)28:08<2311::Aid-immu2311>3.0.Co;2-g (1998).

41 Krieg, C. et al. High-dimensional single-cell analysis predicts response to anti-PD-1 immunotherapy. Nat Med 24, 144–153, doi:10.1038/nm.4466 (2018).

42 De Paoli, P. et al. A subset of gamma delta lymphocytes is increased during HIV-1 infection. Clin Exp Immunol 83, 187–191, doi:10.1111/j.1365-2249.1991.tb05612.x (1991).

43 Behr, M. A., Kaufmann, E., Duffin, J., Edelstein, P. H. & Ramakrishnan, L. Latent Tuberculosis: Two Centuries of Confusion. Am J Respir Crit Care Med, doi:10.1164/rccm.202011-4239PP (2021).

44 McLane, L. M., Abdel-Hakeem, M. S. & Wherry, E. J. CD8 T Cell Exhaustion During Chronic Viral Infection and Cancer. Annu Rev Immunol 37, 457–495, doi:10.1146/annurev-immunol-041015-055318 (2019).

45 Lu, L. L. et al. A Functional Role for Antibodies in Tuberculosis. Cell 167, 433–443 e414, doi:10.1016/j.cell.2016.08.072 (2016).

46 Naluyima, P. et al. Terminal Effector CD8 T Cells Defined by an IKZF2(+)IL-7R(-) Transcriptional Signature Express FcgammaRIIIA, Expand in HIV Infection, and Mediate Potent HIV-Specific Antibody-Dependent Cellular Cytotoxicity. J Immunol 203, 2210–2221, doi:10.4049/jimmunol.1900422 (2019).

47 Klenerman, P. The (gradual) rise of memory inflation. Immunol Rev 283, 99–112, doi:10.1111/imr.12653 (2018).

48 Schober, K. et al. Reverse TCR repertoire evolution toward dominant low-affinity clones during chronic CMV infection. Nat Immunol 21, 434–441, doi:10.1038/s41590-020-0628-2 (2020).

49 Cheng, Y. et al. Non-terminally exhausted tumor-resident memory HBV-specific T cell responses correlate with relapse-free survival in hepatocellular carcinoma. Immunity, doi:10.1016/j.immuni.2021.06.013 (2021).

50 Weber, E. W. et al. Transient rest restores functionality in exhausted CAR-T cells through epigenetic remodeling. Science 372, doi:10.1126/science.aba1786 (2021).

51 Correia, D. V. et al. Differentiation of human peripheral blood Vdelta1+ T cells expressing the natural cytotoxicity receptor NKp30 for recognition of lymphoid leukemia cells. Blood 118, 992–1001, doi:10.1182/blood-2011-02-339135 (2011).

52 Stewart, C. A. et al. Germ-line and rearranged Tcrd transcription distinguish bona fide NK cells and NK-like gammadelta T cells. Eur J Immunol 37, 1442–1452, doi:10.1002/eji.200737354 (2007).

53 Rigau, M. et al. Butyrophilin 2A1 is essential for phosphoantigen reactivity by gammadelta T cells. Science 367, doi:10.1126/science.aay5516 (2020).

54 Vavassori, S. et al. Butyrophilin 3A1 binds phosphorylated antigens and stimulates human gammadelta T cells. Nat Immunol 14, 908–916, doi:10.1038/ni.2665 (2013).

55 Chien, Y. H., Meyer, C. & Bonneville, M. gammadelta T cells: first line of defense and beyond. Annu Rev Immunol 32, 121–155, doi:10.1146/annurev-immunol-032713-120216 (2014).

56 He, X. et al. The potential role of CD16+ Vgamma2Vdelta2 T cell-mediated antibody-dependent cell-mediated cytotoxicity in control of HIV type 1 disease. AIDS Res Hum Retroviruses 29, 1562–1570, doi:10.1089/AID.2013.0111 (2013).

57 Glanville, J. et al. Identifying specificity groups in the T cell receptor repertoire. Nature 547, 94–98, doi:10.1038/nature22976 (2017).

58 Stuart, T. et al. Comprehensive Integration of Single-Cell Data. Cell 177, 1888–1902 e1821, doi:10.1016/j.cell.2019.05.031 (2019).

59 van Dijk, D. et al. Recovering Gene Interactions from Single-Cell Data Using Data Diffusion. Cell 174, 716–729 e727, doi:10.1016/j.cell.2018.05.061 (2018).

60 Wagner, F., Yan, Y. & Yanai, I. K-nearest neighbor smoothing for high-throughput single-cell RNA-Seq data. BioRxiv, doi:10.1101/217737 (2018).

61 Van Gassen, S. et al. FlowSOM: Using self-organizing maps for visualization and interpretation of cytometry data. Cytometry A 87, 636–645, doi:10.1002/cyto.a.22625 (2015).

